# The nature-nurture transform underlying the emergence of reliable cortical representations

**DOI:** 10.1101/2022.11.14.516507

**Authors:** Sigrid Trägenap, David E. Whitney, David Fitzpatrick, Matthias Kaschube

## Abstract

The fundamental structure of cortical networks arises early in development prior to the onset of sensory experience. However, how endogenously generated networks respond to the onset of sensory experience, and how they form mature sensory representations with experience remains unclear. Here we examine this ‘nature-nurture transform’ using *in vivo* calcium imaging in ferret visual cortex. At eye-opening, visual stimulation evokes robust patterns of cortical activity that are highly variable within and across trials, severely limiting stimulus discriminability. Initial evoked responses are distinct from spontaneous activity of the endogenous network. Visual experience drives the development of low-dimensional, reliable representations aligned with spontaneous activity. A computational model shows that alignment of novel visual inputs and recurrent cortical networks can account for the emergence of reliable visual representations.

**One sentence summary:** The reliability of cortical representations emerges from experience-driven reorganization of endogenous networks

---

Cortical circuits emerge from a developmental sequence that includes two distinct phases: an early period prior to the onset of experience during which endogenous mechanisms are thought to formulate the initial framework of cortical networks (*1*–*4*), and a subsequent period during which these early networks are refined under the influence of experience (*5*–*9*). The visual cortex of higher mammals has served as a powerful model for exploring the contributions of these different phases to the development of mature cortical networks. Prior to the onset of visual experience, activity independent mechanisms combine with activity dependent mechanisms driven by patterns of endogenous activity derived from the retina and the LGN (*10*–*12*) to generate a robust modular network structure in visual cortex that is evident in patterns of spontaneous activity (*13*, *14*). This endogenously generated functional network is thought to form the initial framework for the emergent cortical representation of stimulus orientation since visual stimulation at or before eye opening drives weakly orientation-selective responses at the cellular and modular scale (*15*–*19*) and spontaneous activity prior to eye opening is predictive of the representations of stimulus orientations at eye opening (*14*). Despite recognizing the considerable network organization present before experience, we lack a clear understanding of the capacity of the endogenous cortical network to reliably represent stimulus orientation at the onset of visual experience. Currently, we cannot rule out the possibility that the initial responses at eye opening are weak or exhibit only a poor modular structure. Moreover, while patterned visual input appears necessary to improve stimulus selectivity and to align the responses from the two eyes (*15*), the full scope of experience-driven transformations of these initial responses that give rise to mature stimulus representations is unknown at present.

To explore these critical early developmental dynamics, we employed chronic *in vivo* calcium imaging of visually evoked activity in visual cortex of postnatal ferrets prior to and following the onset of visual experience (Fig. 1A; Methods). Utilizing calcium sensors to visualize activity at the modular and cellular scale allowed us to study developing visual representations in the behaviorally relevant regime of individual trials. Our experiments reveal dramatic developmental changes in network response reliability and significant departures of the emerging visual representations from early endogenous network structure that are driven by the visual experience. Based on a computational model whose predictions closely match the biology, we propose that visual experience drives the alignment of feedforward and recurrent networks to transform a nascent modular network with diverse and unreliable visual responses into a mature network with a distinctive modular structure and highly reliable visual responses.

**Figure 1:**
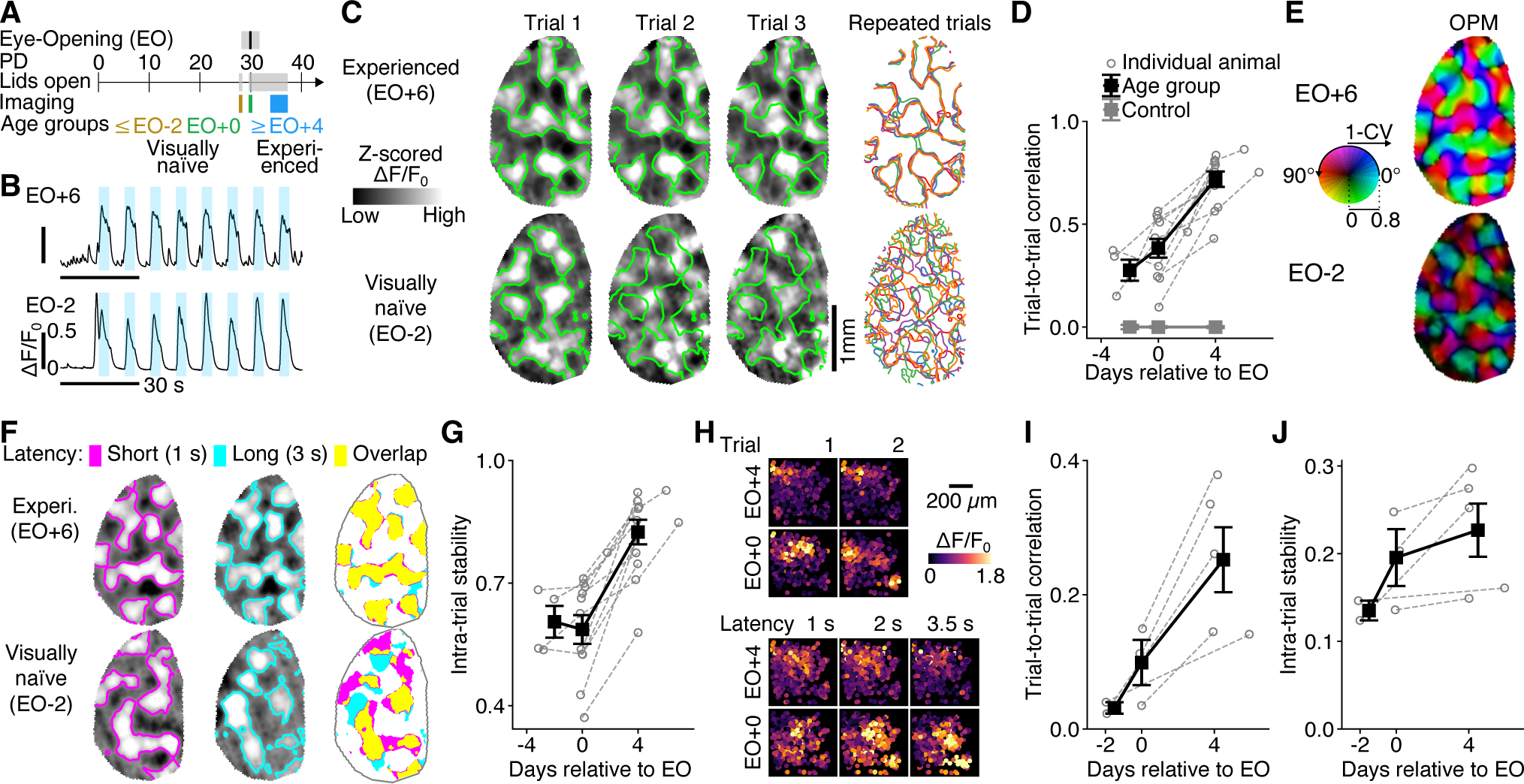
In visually naïve animals, grating stimulus evoked activity is robust and modular, but also highly variable both within and across trials. (A) Timeline of ferret visual cortex development and experiments (PD=postnatal day; EO=eye opening). (B) Grating stimulus evoked activity (GCaMP6s widefield epi-fluorescence imaging), averaged over the region of interest (ROI) in a visually naïve animal, two days prior to eye opening (EO-2; bottom) and in an experienced animal, six days after eye opening (EO+6; top). Light blue: stimulation (moving grating, 16 directions, randomly alternated). (C) Left: Examples of grating-evoked activity patterns for repeated presentations of the same stimulus. Top: experienced (EO+6); bottom: visually naïve (EO-2). Superimposed in green are the zero-contours from the first trial. Activity patterns are taken 0.5 s after stimulus onset. Right: Overlay of contours from n=5 trials (same stimulus). (D) Mean correlation between activity patterns evoked by the same stimulus, averaged over all stimuli (‘trial-to-trial correlation’, see main text and Methods). Animals longitudinally recorded. Error bars show Mean ± SEM for age groups: EO-3 to −2, EO+0, EO+4 to +7 (N= 4,11,11 animals, respectively). Control: random patterns with matched spatial correlation function (Methods). Legend, omitting control, applies also to (G), (I), (J), below. (E) Vector-sum orientation preference map (OPM); angle shown in color and magnitude (1 – circular variance (CV)) in luminance value. (F) Stability of activity pattern within a trial in a visually naïve (EO-2) and an experienced animal (EO+6). (G) Correlation between response patterns 1 s and 3 s after stimulus onset, averaged over all trials and stimuli (‘intra-trial stability’) at different ages. (H) Top: Examples of single cell grating-evoked activity patterns for repeated presentations of the same stimulus, 0.5 s after stimulus onset pooled over 4 imaging planes (GCaMP6 two-photon fluorescence imaging). Bottom: Cellular activity patterns during a trial (same field of view; activity temporally smoothed for illustration using a gaussian kernel, SD=0.33 s). (I) Trial to trial correlation for cellular responses at different ages (N=5 animals). (J) Intra-trial stability of cellular activity, 0.5 s and 2.5 s after stimulus onset at different ages. Same animal presented in (C), (E), (F). For illustration, activity patterns were clipped at 2 SD.

## Strong but highly variable network responses in visually naïve animals

As a frame of reference for the qualities of the earliest visual responses, we start with consideration of the patterns of activity evoked by grating stimuli in animals that have had several days of patterned visual experience (Fig. 1A-C). As expected, single trial responses were strong and modular (comprising a number of distinct active domains) and also highly reliable, such that repeated presentations of the same stimulus evoked highly similar single trial patterns of activity (assessed 0.5 s after stimulus onset, Fig. 1C, upper). Intriguingly, in young, visually naïve animals, with eyes opened prematurely several days prior to the natural time of eye opening (EO), single trial grating evoked activity was also robust (Fig. 1B, lower) and displayed an equally pronounced modular structure (Fig. 1C, lower). However, single trial response patterns were strikingly different for repeated presentations of the same grating stimulus (Fig. 1C, lower).

To quantify this apparent difference in reliability of responses across trials we examined the ‘trial-to-trial correlation’ (see Methods for definition) in multiple animals across ages and found a dramatic increase in trial-to-trial correlation four days following the onset of visual experience (Fig. 1D, EO vs EO+4, p=0.0002, paired permutation test, see Methods), along with a striking increase in orientation discrimination (SI Fig. 1A). At the earliest time points tested, correlations between evoked activity patterns across trials while low, were greater than zero (Fig. 1D; p<10^-5^, random control patterns, see Methods) consistent with the presence of a nascent modular structure at this early stage. Indeed, averaging over trials in visually naïve animals confirmed the presence of a weak but systematic bias in preferred orientation across cortical space (orientation preference map – OPM) (Fig. 1E lower), consistent with previous studies that have found weak structure in trial-averaged activity patterns at these early time points (*15*–*17*, *19*). Our observations suggest that a major contributor to the emergence of mature orientation representations (Fig.1E top) is reduction in response variability at the network scale.

While these results have focused on the visual responses shortly (0.5 s) after stimulus onset, we thought it was important to assess whether a similar degree of instability was present in the dynamics of sustained activity. Our moving visual grating stimuli were presented for a duration of 4-5 seconds, and thus we systematically assessed the degree to which the evoked activity was stably maintained during visual stimulation by comparing the activity pattern 1 s after stimulus onset to that 2 s later. Indeed, following the onset of visual experience, responses were highly stable during the stimulation period (Fig. 1F top), contrasting with responses in the visually naïve cortex that showed a strong tendency to vary during the period of stimulation (Fig. 1F bottom). To quantify this trend, we defined ‘intra-trial stability’ as the correlation between activity patterns 1 s vs. 3 s after stimulus onset averaged over all trials and stimuli (see Methods). We found that across the animals in our sample, intra-trial stability increased significantly several days after eye opening (Fig. 1G; p<0.001, unpaired bootstrap test, see Methods), further emphasizing the high degree of instability in visually evoked responses of naïve animals.

These wide-field imaging results were confirmed with two-photon imaging of the activity patterns in large populations of single neurons. Repeated presentations of the same visual grating stimulus in visually naïve animals evoked single cell responses with amplitude and spatial structure comparable to experienced animals, but with much lower uniformity of responses across trials (Fig. 1H top, SI Fig. 2). Cellular population activity patterns from younger animals were only weakly correlated trial to trial (Fig. 1I), severely limiting orientation discrimination (SI Fig. 1B); but these patterns became progressively more correlated several days after eye opening (Fig. 1I), following a similar trajectory as observed with wide-field imaging in larger fields of view (Fig. 1D). Additionally, population activity patterns in younger animals were not stably maintained during the period of stimulation (Fig. 1H lower, J), consistent with our results using wide-field imaging (Fig. 1G).

Thus, with the initial exposure to activity driven by visual stimuli, endogenously generated circuits deliver responses that are strong and highly structured, but lack the reliability necessary to faithfully represent stimulus orientation. This implies that the weak stimulus tuning that has been observed at the level of trial averages (*15*–*19*) is not due to weak or poorly structured evoked activity, but reflects a significant trial-to-trial variability of strong modular responses. But to shed light on the mechanism that gives rise to highly reliable network responses requires first analyzing the overall organization of the initial grating evoked responses and their relation to the endogenous network structure.

## Reduction in diversity of modular patterns following onset of experience

Our observation of weak consistency of initial responses – both across and within trials – raises the question whether the initial responses to visual grating stimuli consists of a diverse set of patterns, such that the single stimulus parameter visual angle is represented by a higher-dimensional linear manifold in the visually naïve cortex, and potentially by a much more confined manifold in the experienced cortex. To test this, we characterized the overall organization of evoked activity patterns for all trials and stimuli using principal component analysis. The principal components – themselves patterns (SI Fig. 3A) – reveal the most prevalent modes of co-variation between cortical locations across the activity patterns. In experienced animals, the projection of individual (z-scored and normalized) activity patterns into the space spanned by the two leading principal components (SI Fig. 3B) revealed a well-ordered circular structure with radius close to 1, suggesting that stimulus orientation is mapped onto a nearly two-dimensional linear manifold in neural activity space (Fig. 2A, upper). In contrast, we found that in visually naïve animals many projection lengths were much smaller than 1 (Fig. 2A, lower) and the variance of evoked activity was distributed over the principal components more broadly than in experienced animals (Fig. 2B). To quantify this trend, we computed the linear ‘dimensionality’ of grating evoked response patterns (see Methods), which, initially was relatively high, but showed a pronounced decrease with the onset of visual experience (Fig. 2C, SI Fig. 3C). Thus, the early modular responses are more diverse, residing in a higher-dimensional linear manifold than those found in the experienced cortex following the onset of visual experience, suggesting that these initial modular patterns reflect a more flexible repertoire of network responses to visual input, fundamentally different from the experienced cortex.

**Figure 2:**
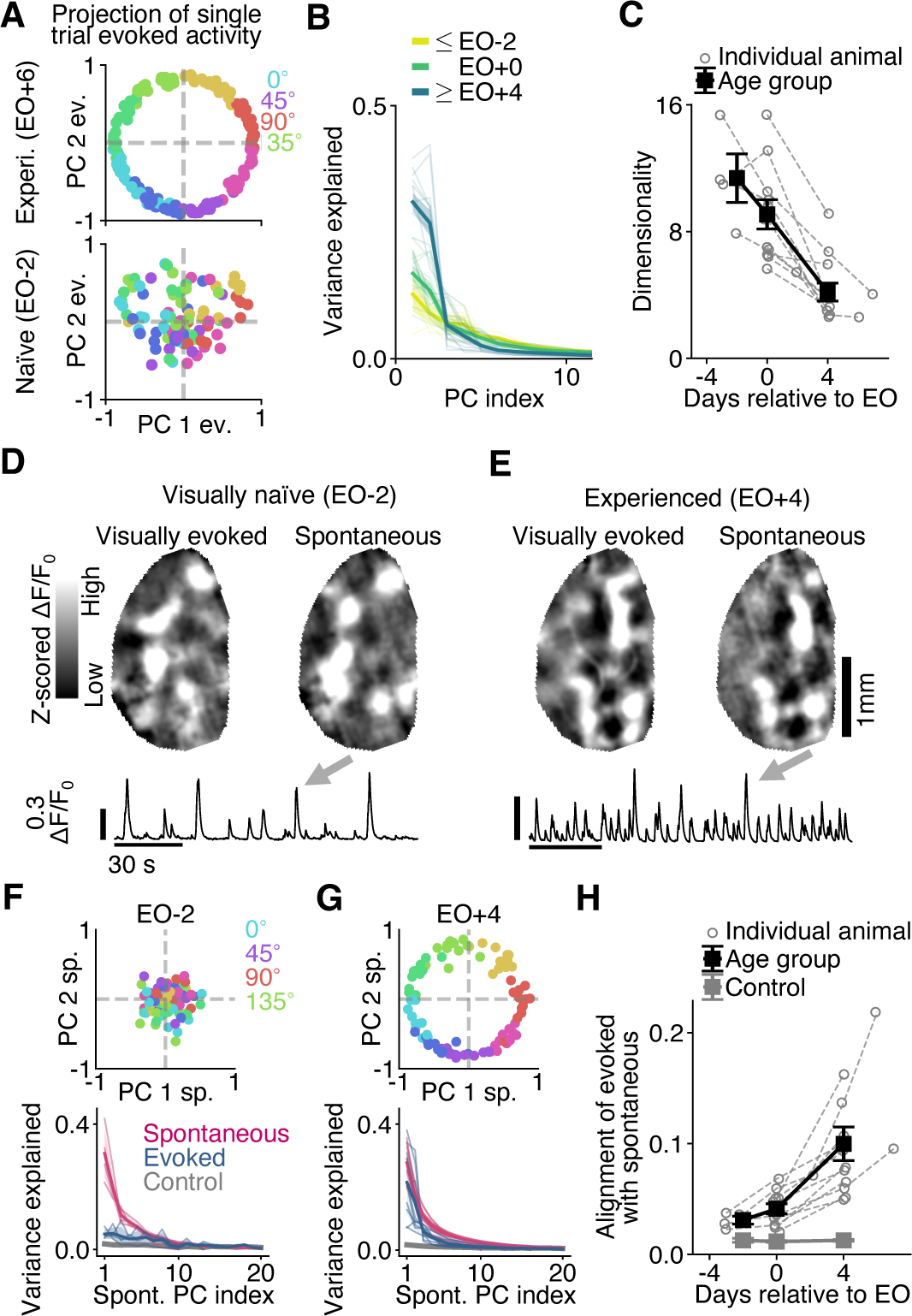
Visually naïve animals show initial grating evoked responses that are diverse and only loosely aligned with spontaneous activity. (A) Single trial grating evoked activity patterns in a visually naïve animal (EO-2, bottom) and in an experienced animal (EO+6, top) projected into the space spanned by the first two principal components (PCs) computed on the respective day. Patterns were represented as vectors over all pixels and normalized to unit length. Here and below, evoked activity is taken 0.5 s after stimulus onset (unless noted otherwise). (B) Variance explained by leading principal components of evoked activity for different age groups (EO-3 to −2, EO+0, EO+4, EO+6: N= 4,11,11,2 animals, respectively). Thick lines, average over animals; thin lines, individual animals. (C) Linear dimensionality (participation ratio, see Methods) of evoked activity. Error bars show Mean ± SEM for age groups: EO-3 to −2, EO+0, EO+4 to +7 (N= 4,11,11 animals, respectively). (D) Spontaneous activity in the visually naïve cortex (EO-2). Bottom: Spontaneous activity trace (black) averaged over the ROI. Top: Spontaneous activity pattern at peak indicated by grey arrow (right), and its best matching (single trial) visual grating evoked activity pattern (left). (E) As (D), but for the (EO+4). (F) Top: Single trial grating evoked activity patterns (normalized to unit length, see Fig. 2B) in the visually naïve cortex projected into the space spanned by the first two PCs of spontaneous activity. Bottom: fraction of variance of spontaneous activity, evoked activity, and control patterns (red, blue, and grey, respectively) explained by the first 20 PCs of spontaneous activity. Thick lines, average over animals (N=11); thin lines, individual animals. Control in (F-H): random patterns with matched spatial correlation function (Methods). (G) As (F), but for the experienced cortex. The number of activity patterns in (C) and (D) were matched across age. (H) ‘Alignment of evoked with spontaneous activity’ defined as the fraction of spontaneous variance explained by an evoked pattern, averaged over all trials and stimuli. Error bars show Mean ± SEM for age groups: EO-3 to −2, EO+0, EO+4 to +7, N= 4,11,11 animals, respectively.

## Initially weak correspondence between evoked and spontaneous activity

Having observed this considerable diversity of early evoked activity patterns, we next sought to shed light on its potential origin by examining the relation of these initial evoked patterns to the endogenously generated network structure, as reflected by the spontaneous activity in the visually naïve cortex. One possibility is that the initial evoked patterns closely correspond to spontaneous patterns of the endogenous network, in which case we would be able to conclude that the patterns that form the initial representations of visual angles already exist in the endogenous network activity. Consistent with this idea, the patterns of spontaneous activity in the early visual cortex were observed to be modular and diverse (*14*), and appear reminiscent of the patterns of evoked activity we observe in visually naïve animals. Moreover, this early modular spontaneous activity is predictive of the columnar organization of the future orientation preference map (*14*) and, different from mice (*20*), a significant structural relation between the two remains evident in the mature visual cortex of carnivores (*14*, *21*, *22*) and primates (*23*). However, it is currently unclear how closely the initial visually evoked responses of the endogenous network correspond to its modular patterns of spontaneous activity.

To test this possibility, we therefore recorded spontaneous activity in the same animals studied above. As reported previously (*14*) we found that patterns of spontaneous activity in the early visual cortex often extended over the entire field of view and were comparable in strength to grating evoked activity (Fig. 2D, E) and exhibited a similar local modular structure (SI Fig. 4). We extracted these patterns at the activity peaks during episodes of 5-10 minutes of spontaneous activity (see Methods) and used principal component analysis to reveal the most prevalent modes of co-variation across spontaneous patterns (see Methods; SI Fig. 3A, D). Overall, the linear dimensionality of spontaneous activity changed only little over the age range considered (SI Fig. 3D-F). Exploring the relationship between grating evoked and spontaneous activity recorded on the same day, we observed an only weak correspondence between the two in visually naïve animals, while their alignment became tight only following visual experience (Fig. 2F, G). Projecting single trial evoked activity patterns onto the two leading principal components of spontaneous activity revealed in experienced animals a representation of the cyclic stimulus space that was comparable in the degree of order to the projections onto the two leading evoked components at the same age (Fig. 2G top; compare Fig. 2A top). Projection lengths were close to 1 and, consistently, most visually evoked variance was concentrated on the two leading principal components of spontaneous activity (Fig. 2G bottom; see SI Fig. 5E-G for similar results in awake animals). In contrast, in visually naïve animals two days prior to eye opening, projection lengths of evoked patterns onto leading spontaneous components were consistently smaller than 1 (Fig. 2F top) and the overlap onto the main spontaneous components, while still noticeably larger than for random control patterns (Fig. 2F bottom; p<0.001, see Methods), was considerably smaller than the values found when projecting the evoked onto the main evoked components (Fig. 2B). Defining ‘alignment of evoked with spontaneous’ as the fraction of total spontaneous variance that an individual evoked pattern on average explains, we observed a several-fold increase after eye opening (Fig. 2H, p=0.0003, paired permutation test, see Methods; consistent with alternative alignment measures, SI Fig. 5A-D). Such trend is reminiscent of reports at a later developmental stage in ferrets (*24*), while different from the trend observed in zebrafish (*25*). Our results thus argue against the possibility that the patterns of evoked activity, at the onset of vision, closely correspond to patterns of the endogenous network activity, and argue that these initial evoked responses consist of novel patterns. Moreover, while only loosely aligned at eye opening, single trial evoked patterns strongly align with spontaneous patterns following visual experience.

## Normal patterned visual experience is necessary to achieve mature network properties

Having seen the transformation from visual responses that are variable and only loosely aligned with spontaneous activity in the visually naïve cortex to the reliable and well aligned responses several days later, we next sought to test the degree to which visual experience drives these developmental changes by studying responses in binocularly deprived animals. These animals (N=3) were lid-sutured for one week starting at P30 (the normal time of eye opening), preventing visual stimulation through the open eyes prior to the experiments (at around P38) (Fig. 3A; see Methods). As before, we recorded both visual grating evoked and spontaneous patterns of activity in all animals. Binocular deprivation had a significant impact on all of the four cortical network properties of grating evoked activity that we have shown change substantially after the onset of visual experience: trial-to-trial correlation, intra-trial stability, dimensionality and alignment with spontaneous activity. Principal component analysis revealed that evoked responses lacked the orderly circular distribution in two dimensions that is evident in normally reared animals and age-matched controls (Fig. 3B; compare Fig. 2A upper). Indeed, visual responses in visually deprived animals lacked the reliability found in age-matched, visually experienced controls, both in trial-to-trial correlation and intra-trial stability (Fig. 3C, D; BD versus Control, c: p=0.017, d: p=0.021, unpaired bootstrap test) and showed weak orientation discrimination (SI Fig. 1C). Likewise, with deprivation, the diversity of visually evoked patterns was not significantly different from that found in visually naïve animals, with no sign of the decrease in dimensionality found in animals with experience (Fig. 3E, BD versus EO, p=0.38, unpaired bootstrap test). Finally, in the deprived animals, projecting evoked activity patterns onto the two leading principal components of spontaneous activity did not reveal an orderly structure (Fig. 3F; compare Fig. 3C top) and the overlap with the main spontaneous components, while noticeable, remained near the level of visually naïve animals (Fig. 3G; compare Fig. 2G bottom). Quantitatively, we did not observe any increase in alignment between evoked and spontaneous activity in deprived animals beyond what was observed in normal animals at the time of eye opening (p=0.971, unpaired bootstrap test; Fig. 3H). These results indicate that normal visual experience is critical for changes in a number of cortical network properties that accompany the emergence of reliable visual representations.

**Figure 3:**
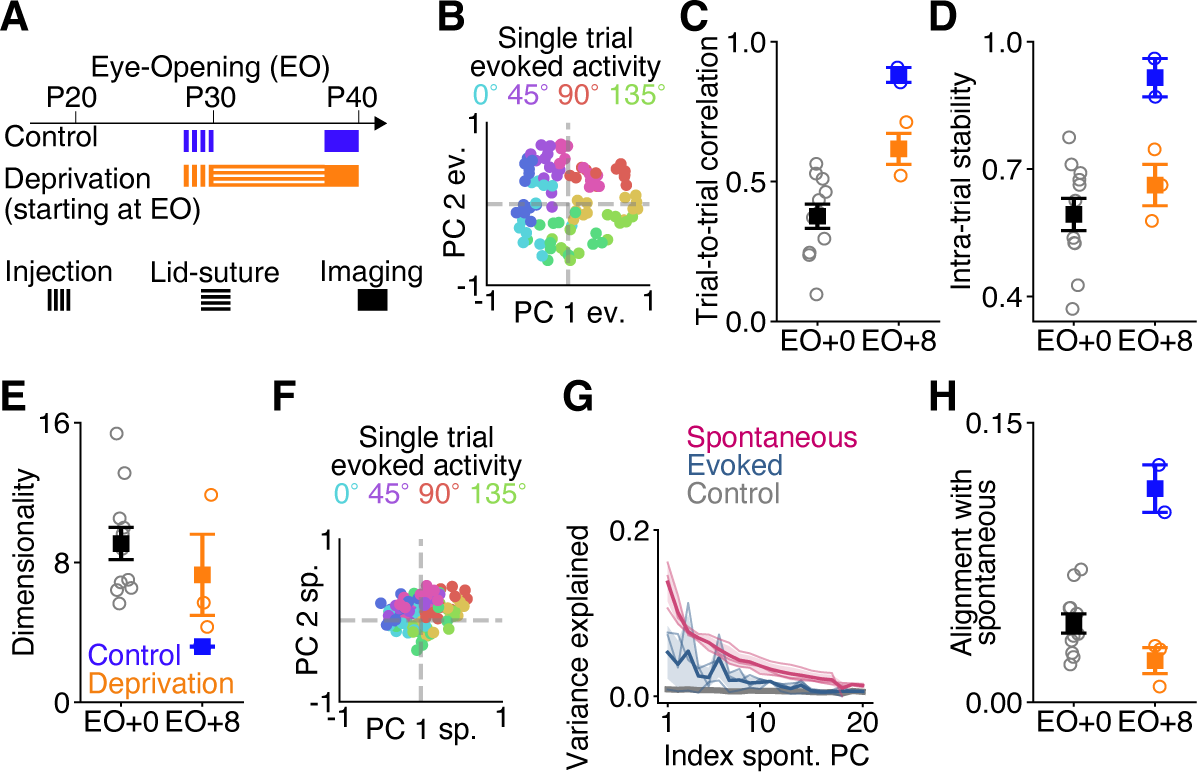
Patterned visual experience is required to develop low-dimensional, stable visual responses aligned with the spontaneous network activity. (A) Experimental timeline for deprivation (orange) and control (blue) experiments. (B) Evoked activity in deprived animals projected into the space spanned by the first two PCs of evoked activity computed on the same day (8 days after the normal time of eye opening). Activity was normalized to unit length (compare Fig. 2A). (C) Trial-to-trial correlation in deprived (N=3, orange) and control (N=2, blue) animals, in comparison to visually naïve animals at EO+0 (black, reproduced from Fig. 1D). Circles: single animals; error bars: Mean ± SEM. (D) As (C), but for intra-trial stability (naïve animals reproduced from Fig. 1G). (E) As (D), but for linear dimensionality (naïve animals reproduced from Fig. 2C). (F) Same as (B), but projected into the spontaneous PC space. (G) Fraction of variance of spontaneous activity, evoked activity, and control patterns (red, blue and grey, respectively) explained by the first 20 PCs of spontaneous activity for deprived animals. (H) As (C), but for alignment of evoked with spontaneous activity (naïve animals reproduced from Fig. 2H).

## The feedforward-recurrent alignment hypothesis

While, in principle, these network properties could change independently from one another (SI Fig. 6), the fact that they mature over the same period of time, and depend on experience, made us wonder whether they could be the product of a common underlying mechanism that builds reliable network responses. Ample computational work suggests that recurrent connections can give rise to amplification within subnetworks of coactive network units (*26*–*31*). Input that aligns more with such a subnetwork is expected to elicit a more robust response – an amplification that reflects greater resonance with the intrinsic network structure (Fig. 4A). At eye opening, the cortex is exposed to structured visually driven input for the first time and, assuming the two are, initially, not properly aligned, we reasoned that proper alignment of this novel feedforward drive to the cortical recurrent network could be an important factor in the maturation of faithful network responses to visual input.

**Figure 4:**
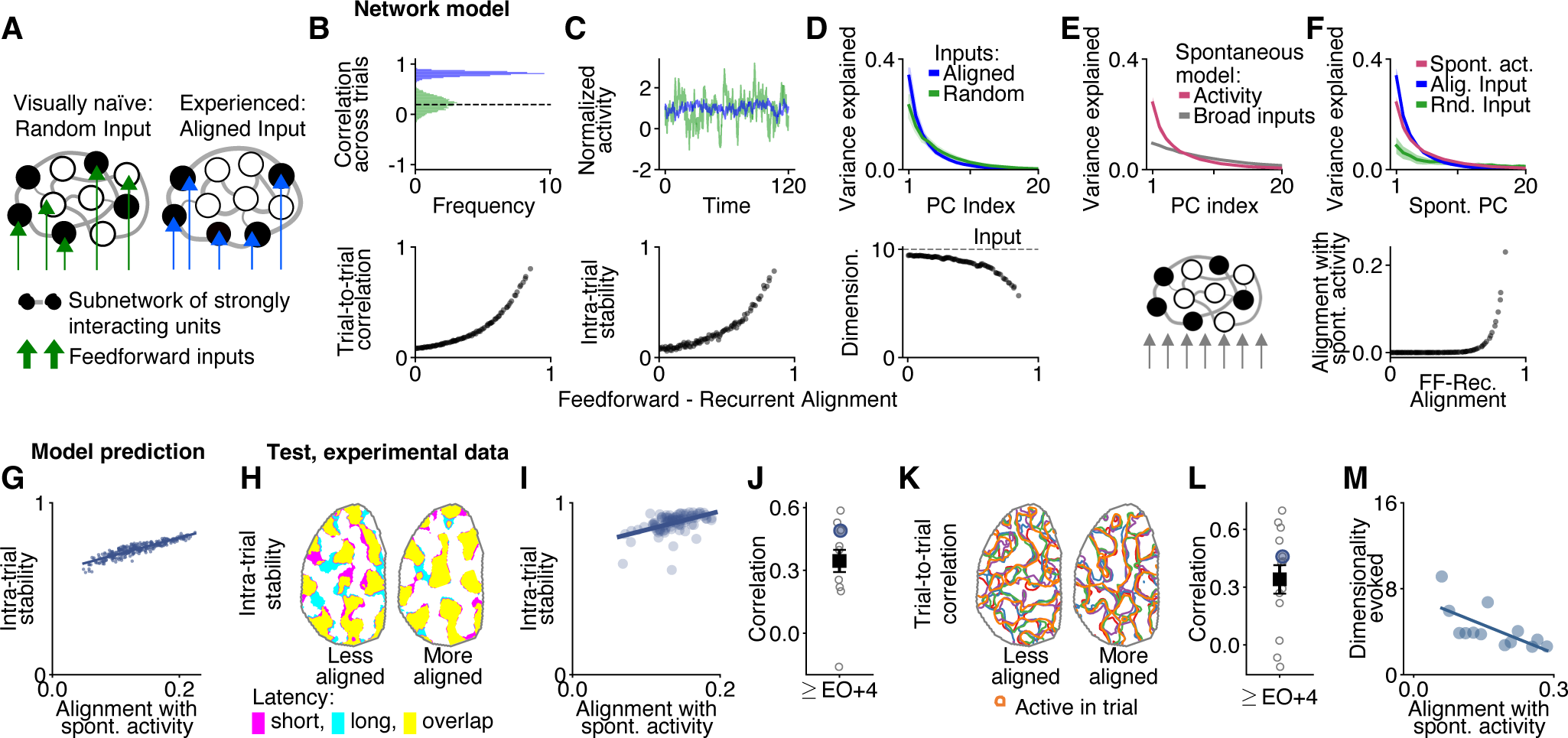
Feedforward-recurrent alignment can account for the observed changes in network activity properties. (A) Schematic illustration of the feedforward-recurrent alignment hypothesis: In the experienced cortex, the feedforward input is aligned and ‘resonates’ with the recurrent network (right), whereas in the naïve cortex their relationship is random (left). (B-F) Computational model. (B) Top: Trial-to-trial correlation for random input (green) and input maximally aligned to the recurrent network (blue). Variability across trials is modelled by adding Gaussian noise (zero mean, unit variance) to each input (see Methods). Bottom: Trial-to-trial correlation as a function of feedforward-recurrent alignment. (C) As (B), but for intra-trial stability. Variability within trials is modelled by adding white Gaussian noise to each input starting from the steady-state evoked response (see Methods). Top shows activity of a network unit (normalized by average response). (D) As (B), but for dimensionality. Top shows variance explained by leading PCs of evoked activity (see Fig. 2B,C for comparison to experiment). Input is a 10-dimensional Gaussian distribution (see Methods). (E) Model of spontaneous activity (schematic, bottom): A broad, high-dimensional distribution of inputs aligned with the recurrent network (see Methods), producing a lower-dimensional output (top). (F) Alignment of evoked to spontaneous activity (see Fig. 2F,G (bottom) for comparison to experiment). Shaded region: SEM around mean (N=50 networks). (G) Model prediction: trials for which the steady-state evoked response is more aligned to spontaneous activity also show higher intra-trial stability. N=300 samples drawn for the input distribution used in (D) with optimal alignment and temporally varying noise as in (C). (H-J) Test, experimental data: (H) Outline of grating evoked activity 1 s (magenta) and 3 s (cyan) after stimulus onset for a response pattern more weakly aligned (left) and for one more strongly aligned (right) to spontaneous activity 0.5 s after stimulus onset. Intra-trial stability: 0.63 (left), 0.90 (right); alignment to spontaneous activity: 0.06 (left), 0.18 (right). Day: EO+4. Same animal as in Fig. 1F. (I) As (G) but for experimental data using all trials for all stimuli from the animal shown in (G) at EO+4. (J) Correlation for the data shown in (I) (large blue circle) and for all other animals in the oldest age group (grey circles; 13 experiments from N=11 animals at EO+4 to +7). Error bars show Mean ± SEM. respectively). (K) Contours show outlines of grating evoked patterns (taken 0.5 s after stimulus onset) from different trials for a stimulus for which the average pattern is less (left) or more (right) aligned to spontaneous activity. Trial-to-trial correlation: 0.75 (left), 0.84 (right); alignment to spontaneous activity: 0.13 (left), 0.18 (right). Same animal as in Fig. 1C. (L) As in (J), but quantifying the relationship illustrated in (K). (M) Evoked dimensionality co-varies with the alignment of the first evoked PC to spontaneous activity (see Methods). 13 experiments from N=11 animals at EO+4 to +7 Data presented in (H), (I) and (K) is from the same animal

To explore this possibility, which we call the ‘feedforward-recurrent alignment hypothesis’, we studied a conceptual model of early cortex and its response to visually evoked input using a minimal linear recurrent network of rate units (see Methods). Each unit represents the pooled activity in a local group of neurons. Connections between units describe the net interactions between local pools. These interactions can be positive or negative and are assumed to be symmetric, for simplicity, with random values drawn from a Gaussian distribution. Importantly, the interactions are assumed to be strong enough to influence activity patterns, consistent with the observation that spontaneous cortical activity exhibits a modular structure prior to eye opening, even after silencing LGN (*14*). The network receives stimulus-driven, feedforward input that varies across neural populations, reflecting random biases in their input selectivity.

We asked whether differences in the degree of feedforward-recurrent alignment (while keeping the statistical properties of the feedforward input and of the recurrent network unchanged) could reproduce the characteristics of network behavior that distinguish naïve and experienced V1 evoked responses. We computed the steady state responses of the network to a static input pattern to which we added a standard normal random vector to model trial-to-trial variability (Methods). Similarly, to model within-trial variability, we added standard normal white noise to the input and computed the correlation of responses at two different time points. Constructing input patterns with varying degree of alignment to the network (see Methods) revealed that those inputs that align better tend to give rise to less variable network responses, and this trend is systematic both across trials and within a trial (Fig. 4B, C). The reason for this trend is the stronger ability for aligned input to drive network responses, relative to the noise component, whose contribution is independent of alignment within this linear framework. Moreover, to reveal how the feedforward-recurrent alignment affects the dimensionality of the activity evoked by a low-dimensional stimulus set, such as moving visual gratings, we considered the generic model of static input being sampled from a low-dimensional (correlated) Gaussian distribution with predefined degree of alignment (see Methods), observing that the dimensionality of the evoked activity patterns is smaller the better-aligned the input distribution is to the network (Fig. 4D).

Moreover, this feedforward-recurrent alignment framework also provides a way for conceptualizing patterns of spontaneous activity and their relation to evoked activity in naïve and experienced animals. We assume that spontaneous activity in the visual cortex results from a broad range of inputs involving various sources, and that these modular patterns, driven by endogenous activity, reflect the current state of aligned feedforward-recurrent network. Robust modular spontaneous activity patterns are present at least 10 days prior to eye opening (*14*) indicative of a feedforward-recurrent network alignment conditioned solely by endogenous patterns of activity in visually naïve animals. Modeling these endogenous inputs as samples drawn from a high-dimensional Gaussian distribution (Fig. 4E, bottom), the distribution of network activity patterns resulting from these endogenous inputs transformed via the recurrent network, is much lower-dimensional than the input distribution (Fig. 4E, top; Methods). Assuming at eye opening visual experience drives novel patterns of feedforward input that are not as well aligned to the recurrent network as the endogenous patterns, the evoked and spontaneous pattern distributions overlap only little (Fig. 4F, top, green vs. red line), similar to what we observed in visually naïve animals (Fig. 2F). Experience-driven changes that optimize the feedforward-recurrent alignment would then result in distributions of evoked and spontaneous patterns of activity that strongly overlap (Fig. 4F, top, blue vs. red line), consistent with our observation in the visually experienced cortex (Fig. 2G). Thus, in this generic, conceptual model, all four properties – trial-to-trial correlation, intra-trial stability, dimensionality of evoked activity and its alignment to spontaneous activity – are intricately linked to the feedforward-recurrent alignment, suggesting that the immaturity of these properties that we observe in the visually naïve cortex arises from a lack of alignment of feedforward and recurrent networks, while improving this alignment for visually driven input underlies, in part, the changes in the properties we observe after the onset of visual experience.

Finally, this led us to ask whether our biological data is consistent with key predictions of this feedforward-recurrent alignment model. In the model, these four properties are linked due to the common underlying mechanism: network amplification of inputs aligned to the structure of recurrent interactions. Notably, for input well aligned to the recurrent network this model predicts a tight link even at the level of individual trials, namely that responses that exhibit greater similarity with the leading components of spontaneous activity also show higher intra-trial stability (Fig. 4G). Therefore, if this mechanism also holds for the cortex, then we expect to see such a trend also in the visually experienced age group, for which we assume a strong alignment to the recurrent network. Indeed, when analyzing the responses for all trials of all stimuli for animals in this age group, we observed that the larger the alignment of an individual evoked response pattern with spontaneous activity 0.5 s after stimulus onset, the more stable this response tended to be during the following few seconds of stimulation (Fig. 4H, I). Moreover, for trial-to-trial correlation we found an analogous trend (Fig. 4J, K; SI Fig. 7A-F) and, likewise, for dimensionality, when comparing mean values across different animals (Fig. 4M, SI Fig. 7E). Together, these results show that these tight links between network properties that the proposed mechanism implies is indeed consistent with our empirical data.

## Discussion

Our results demonstrate that experience plays a critical role in transforming an endogenously structured modular network with robust, but diverse and unreliable visual responses into a mature network with a distinctive modular structure and highly reliable visual responses. Following the first steps in the development of visual representations with single trial resolution, we find that what appears to be an increase of initially weak stimulus tuning when observed at the level of trial averages (*15*–*19*), is better described as an increase in consistency both across and within trials of surprisingly robust modular visual responses. Moreover, we find that the initial evoked responses at eye opening consist of novel patterns, distinct from spontaneous activity of the endogenous network and that visual experience drives the development of low-dimensional, reliable representations aligned with spontaneous activity.

Based on our computational model, we propose that these transformations involve the alignment of feedforward and recurrent networks in a fashion that is optimal for amplifying the novel activity patterns induced by the onset of visual experience. Such alignment is a property of the larger network, ensuring that individual activity domains act in concert when driven by visual input. Achieving alignment could involve changes in the feed-forward input to the recurrent network, changes in the recurrent network, or some combination of the two. We note that our model does not specify changes in the selectivity of feedforward inputs that are typically regarded as critical for development of the representation of orientation (*29*, *32*–*37*). Nevertheless, the process of feedforward-recurrent network alignment could pave the way for the emergence of highly-selective feedforward responses following the onset of visual experience. The specific neural circuit changes that produce feedforward and recurrent network alignment and the plasticity mechanisms that enable them remain to be determined. Our observation that grating evoked activity, when falling closer to the main components of spontaneous activity, is more stable both within and across trials is consistent with a process of dynamic self-organization (*38*) in which more aligned, stable and thus more correlated activity patterns are strengthened by mechanisms of activity dependent synaptic plasticity. It remains for future studies to determine the exact sequence of developmental changes that drives alignment of feedforward and recurrent circuits enabling the emergence of highly reliable stimulus representations.

## Acknowledgements

We would like to thank J. Drayer and D. Ouimet for technical assistance, as well as members of the Fitzpatrick and Kaschube laboratories for helpful discussions. We thank Gordon B. Smith and Bettina Hein for early support of the project and valuable input. We thank Jeremy T. Chang for technical assistance and sharing datasets. This research was supported by US National Institutes of Health grants EY011488 and EY026273 (D.F.), BMBF project D-USA-Verbund: SpontVision, FKZ 01GQ1507 (M .K.) and ProLoewe CMMS (M.K, S.T.), the International Max Planck Research School for Neural Circuits in Frankfurt (S.T.), as well as the Max Planck Florida Institute for Neuroscience.

## Author contributions

All authors designed the study, analyzed the results, and wrote the paper. D.E.W. performed the wide-field and 2-photon calcium imaging, S.T. and M.K. performed the computational modelling.

## Author information

The authors declare no competing financial interests. Correspondence and requests for materials should be addressed to D.F. or M.K.

## Code availability

The code for data analysis is available from the corresponding author upon request. The code for the model will be made available on GitHub upon publication.

## Data availability

The data that support the findings of this study are available from the corresponding author upon reasonable request.

## Methods

### 1 Experiments

The experimental procedures follow (*1*) and (*2*) and are described in this section. All experimental procedures were approved by the Max Planck Florida Institute for Neuroscience Institutional Animal Care and Use committee and were performed in accordance with guidelines from the U.S. National Institute of Health. Juvenile female ferrets from Marshal Farms were co-housed with jills on a 16 h light/8 h dark cycle. Parts of the raw data used in this study were used to address different scientific questions in a previous publication (*1*). Also, the results presented in Fig. SI 5E-G are based on raw data used to address different scientific questions in a previous publication (*2*)

### 1.1 Viral injections

Viral injections were performed as previously described (*1*)(*2*). Briefly we expressed GCaMP6s by microinjection of AAV2/1-hSyn-GCaMP6s-WPRE-SV40 (University of Pennsylvania Vector Core) into the visual cortex 6-14 days before imaging experiments. In developing ferrets, viral expression using the hSyn promoter has previously been demonstrated to primarily label excitatory neurons (*3*) and yield multiple millimeters of roughly uniform labeling of GCaMP6s (*2*). Anesthesia induction was performed using either ketamine (50 mg/kg, IM) and/or isoflurane (1%–3%) delivered in O2 and then maintained with isoflu-orane (1%–2%). Atropine (0.2 mg/kg, IM) was administered to reduce secretions, while Buprenorphine (0.01 mg/kg, IM) and a 1:1 mixture of lidocaine and bupivacaine (injected directly into the scalp) were administered as analgesics. Animal temperatures were maintained at 37°C using a homeothermic heating blanket. Animals were also mechanically ventilated and both heart rate and end-tidal CO2 were moni-tored throughout the surgery. Under aseptic surgical technique, a small craniotomy was made over visual cortex 6.5-7 mm lateral and 2 mm anterior to lambda. Approximately 1 mL of virus was pressure infused into the cortex through a pulled glass pipette across two depths (200 mm and 400 mm below the surface). This procedure reliably produced robust and widespread GCaMP6s expression in excitatory neurons over an area > 3 mm in diameter (*2*). To improve the uniformity of the GCaMP6s expression, sometimes an additional injection of 1 mL of virus was pressure injected into a separate region of cortex displaced 1-1.5 mm away from the first injection site.

### 1.2 Eyelid suture procedure

In deprivation and chronic imaging experiments, eyelids were binocularly sutured during a short surgical procedure between P26-30. Anesthesia was induced with isoflurane (2%–5%) and Buprenorphine (0.01 mg/kg, IM) was administered. Using aseptic technique, both eyelids were sutured shut using continuous sutures (6-0 Ethilon suture). Eyelid sutures were monitored daily until removed during an imaging experiment.

### 1.3 Cranial window surgery

All animals were anesthetized and prepared for surgery as described above. In acute imaging experiments, skin and muscle overlying visual cortex were reflected and a custom-designed metal headplate (8 mm DIA) was implanted over the injected region with MetaBond (Parkell Inc.). Then a craniotomy and a subsequent durotomy were performed, and the underlying brain stabilized with a custom-designed titanium metal cannula (4.5 mm DIA, 1.5 mm height) adhered to a thin 4 mm coverslip (Warner Instruments) with optical glue (71, Norland Products, Inc). The headplate was hermetically sealed with a stainless-steel retaining ring (5/16” internal retaining ring, McMaster-Carr) and glue (VetBond, 3M or Krazy Glue).

To allow repeated access to the same imaging field in chronic imaging experiments, we implanted a cranial windows in each animal 2 days prior to the first imaging session. Using aseptic surgical technique, we adhered a metal headpost (7×7 mm) to the skull 3.5 mm anterior of bregma and a separate custom-designed, chamber implant overlying the injected region using MetaBond and black dental acrylic. At the end of the survival cranial window implant surgery, the metal cannula was sealed with a silicone polymer plug (Kwik-kast, World Precision Instruments) to protect the imaging window between imaging experiments. Whenever the imaging quality of the chronic cranial window was found to be suboptimal for imaging, the chamber was opened under aseptic conditions, any regrown tissue/neomembrane was removed, and a new coverslip was inserted.

### 1.4 Imaging experiments

Acute imaging experiments began immediately after the cranial window surgery. For survival imaging experiments, where animals had a cranial window implanted days earlier, anesthesia was induced with isoflurane (2%–5%) and atropine (0.2 mg/kg) was administered. Animals were intubated and ventilated, and an IV catheter was placed in the cephalic vein. The silicon polymer plug overlying the sealed imaging chamber was gently peeled off.

For both acute and survival imaging experiments, eyelid sutures were removed or eyelids were separated where applicable to ensure visual stimulation was always presented to open eyes. Phenylephrine (1.25%–5%) and tropicamide (0.5%) were applied to the eyes to retract the nictitating membrane and dilate the pupil, and the cornea was protected with regular application of eye drops (Systane Ultra, Alcon Laboratories). Prior to imaging, isoflurane levels were reduced from a surgical plane to 1%–1.5%. After reaching a stable, anesthetic baseline for 30 minutes (280-300 bpm), animals were paralyzed with pancuronium bromide (0.1 mg/kg/hr in lactated Ringer’s with 5% dextrose, delivered IV). Upon completion of imaging in acute experiments, isoflurane was raised to 5% and given Euthasol (0.5 ml, IV). In survival experiments, animals were instead recovered from anesthesia and returned to their home cages. During recovery, up to 3 repeated doses of neostigmine (0.06 mg/kg/hr, IV) and atropine (0.05 mg/kg/hr, IV) were used to reverse paralysis.

### 1.5 Calcium signal measurement

Widefield epifluoresence imaging of GCaMP6s was achieved with a Zyla 5.5 sCMOS camera (Andor) controlled by mManager (*4*). Images were acquired at 15Hz with 4×4 binning to yield 640×540 pixels through a 4x air-immersion objective (Olympus, UPlanFL 4x N/0.13NA). For analysis, images were spatially downsampled by a factor of 2× to yield 320×270 pixels at a spatial resolution of 11.63 mm/pixel. The region of interest (ROI) was manually selected to include all parts of the field of view showing strong GCaMP6s expression and a robust visually evoked signal.

Two-photon imaging of GCaMP6s was performed with a B-Scope microscope (ThorLabs) driven by a Mai-Tai DeepSee laser (Spectra Physics) at 940 nm. The B-Scope microscope was controlled by Scan-Image (*5*) in a resonant-galvo configuration with multi-plane images (512×512 pixels) being sequentially collected across either one or four imaging planes using a piezoelectric actuator for an effective frame rate of 30 Hz or 6 Hz respectively. Images were acquired at 2x zoom through a 16x water immersion objective (Nikon, LWD 16X W/0.80NA) yielding cellular fields-of-views of 0.7mm on each side (1.36 mm/pixel). The location of the two-photon calcium imaging ROIs was primarily determined by finding a location with strong GCaMP6s expression and not occluded by large superficial vessels.

### 1.6 Visual stimulation

Visual stimuli were delivered on a LCD screen placed approximately 25–30 cm in front of the eyes using PsychoPy (*6*). To evoke orientation responses, full-field square gratings at 100% contrast, at 0.015 or 0.06 cycles per degree and drifting at 1 or 4 Hz were presented at 16 directions. For optimal responsivity, a single paired spatial and temporal frequency was used for each animal to assess binocular orientation-specific responses. Stimuli were randomly interleaved and presented for 4 s followed by 3-6 s of gray screen. Spontaneous activity was recorded in a darkened room, with the visual stimulus set to a black screen.

## 2 Data analysis

All data analysis was performed using custom written scripts in either Python, MATLAB, or ImageJ.

### 2.1 Preprocessing for widefield imaging data

Following (*2*), all fluorescence images recorded on the same day were corrected for in-plane motion by aligning them (rigid alignment in x and y position) to the preceding frame via maximal 2D cross correlation. This alignment was primarily based on the high contrast vascular structure present in the raw images.

For further analysis, pixels contaminated by blood-vessel signals were removed from the images by generating a mask covering the vascular structure and setting the pixels within this mask to the values outside the ROI. The remaining pixels defined a new ROI, which we used for all analyses. The vascular mask was generated by high-pass filtering the time-averaged fluorescence image and selecting all pixels with a fluorescence smaller than *µ*_*F*_ − 2*σ*_*F*_, where *µ* and *σ* are the mean and standard deviation, respectively, of this time-averaged fluorescence image across space. Overall, we found that similar results were obtained when including the signal from the pixels in the vascular mask. In all patterns depicted in the figures the vascular mask pixels were not removed, for illustration purposes.

The baseline F0 for each pixel was obtained by applying a moving rank-order filter to the raw fluorescence trace (25-30th percentile) with a time window of 60 s. The baseline corrected evoked activity was calculated as (F-F0)/F0 = DF/F0.

#### Stimulus evoked responses

Visual grating evoked responses at time t after stimulus onset were calculated as the average DF/F0 over the period (*t*−0.066 ms, *t*+0.066 ms). Trial-averaged responses at time t were calculated by taking the average across repeated trials to the same stimulus direction at time t after stimulus onset. For all analyses, we used *t* =0.5 s, apart from the analysis of intra-trial stability, in which case we compared the response between *t* =1.0 s and *t*=3.0 s after stimulus onset (see below).

#### Patterns of spontaneous activity

To extract the large-scale patterns of spontaneous activity, we followed the approach described in (*2*). Briefly, we first determined active pixels on each frame using a pixel-wise threshold set to 4 SD above a pixel’s mean value across time, where the SD was estimated from the distribution obtained from the pixel’s activity values below the mean and mean-reflected copies of these. A frame was named active if >80% of its pixels within the ROI were active. Consecutive active frames were combined into a single event starting with the first high-activity frame and then either ending with the last high-activity frame or, if present, an activity frame defining a local minimum in the fluorescence activity. To assess the spatial pattern of an event, we extracted the maximally active frame for each event, defined as the frame with the highest activity averaged across the ROI. Imaging sessions in which fewer than ten spontaneous events were detected were excluded from further analysis (*2*). Typically, the number of events per unit imaging time increased with age and allowed us to gather >30 spontaneous patterns around eye opening and >75 patterns 4-6 days after eye opening.

#### Spatial filtering

For analysis of the wide-field epifluorescence imaging data, we applied spatial band-pass filtering following (*2*) to reduce spatial inhomogeneities in signal strength and photon noise. Briefly, we downsampled the DF/F0 images to 160 × 135 pixels and spatially filtered them using two-dimensional Gaussian filter kernels. For high-pass filtering, we subtracted from each downsampled DF/F0 image a smoothed version of it obtained by applying a Gaussian filter with SD *s*_high_ = 196 *µ*m and normalizing the filtered values by the overlap of the filter kernel with the ROI mask, to correct for boundary effects. Subsequently, we smoothed this image with a Gaussian filter kernel with SD *s*_low_ = 26 *µ*m for low pass filtering. The choice of these filter values largely preserved the modular activity structure of grating evoked and spontaneous activity in ferret visual cortex (*2*).

### 2.2 Preprocessing for two-photon imaging data

#### Cellular ROIs

Cellular ROIs were manually drawn using custom software in ImageJ (Cell Magic Wand) (*3*). The baseline F0 for each pixel/cell was obtained by applying a moving rank-order filter to the raw fluorescence trace (25-30th percentile) with a time window of 60 s. The baseline corrected evoked activity was calculated as (F-F0)/F0 = DF/F0.

#### Morphing across age

To locate the same field-of-views between chronic imaging sessions, we acquired a series of z stack images across different depths incremented at 5 *µ*m steps during each experiment. For each chronic experiment, the original field-of-view was carefully compared to the different z-locations, and a matched location was chosen based on the relative similarity of physical landmarks, such as penetrating blood vessels. We only chose cells for analysis that were common to both sessions. To aid the experimenter in selecting common cells, all fields-of-view were registered into a common reference frame using a 2D elastic deformation (BUnwarpJ plug-in in ImageJ). To determine whether a pair of cell ROIs across chronic experiments corresponded to the same cell, we putatively required that the centroid of the candidate cell pairs were located within 4-5 *µ*m and overlapped more than 25%. The experimenter then manually decided whether the match was reasonable based on similarity of the local neighborhood and physical shape of the cell ROIs.

#### Stimulus evoked activity

Visual grating evoked responses at time t after stimulus onset (Fig 1H-J) were calculated as the average DF/F0 over the period (*t*-0.166 ms, *t*+0.166 ms). We used *t* = 0.5 s, apart from intra-trial stability, for which we compared the response between *t* = 0.5 s and *t* = 2.5 s after stimulus onset (see below). For Figs. SI 1 and SI 2 grating evoked responses were averaged over the full stimulus period.

### 2.3 Quantification

#### Overlap between activity spaces

In several instances (see below), we were interested in the degree to which the activity patterns of a set *A* reside in the most prevalent linear dimensions of a second set of activity patterns *B.* To estimate this quantity, we computed the fraction of variance of patterns *A* that is explained by the leading principal components of the patterns in *B.* Concretely, the variance of *A* explained by the *i*-th principal component **p**_*i,B*_ is

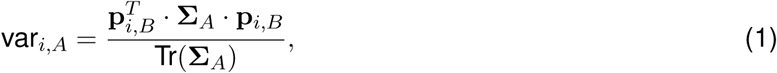

where **Σ**_*A*_ is the covariance matrix of the pattern set *A* and the principal component vectors **p**_*i,B*_ are normalized to unit length. The principal components were estimated using sklearn (*7*) with the LAPACK solver.

#### Trial-to-trial correlation

To measure how reliably the visual cortex responds to repeated presentations of an identical moving grating stimulus (Fig. 1C-D, I), we first computed the correlations (here and below we used the Pearson’s correlation coefficient, if not noted otherwise) between all possible pairs of single-trial response patterns for that stimulus and took the mean over these correlations:

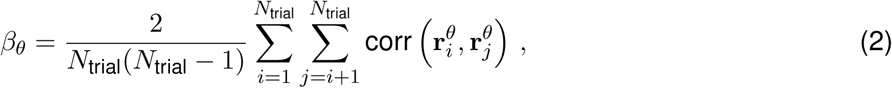

where *N*_trial_ are the number of trials and r_i_^*θ*^ is the response pattern for the *i*-th trial of stimulus *θ* (taken at *t* = 0.5 s after stimulus onset). This mean correlation was then averaged over the *N*_stim_. moving grating directions *θ* to define the ’trial-to-trial correlation’ of responses for an animal on a given day.

#### Orientation Tuning

Orientation tuning for widefield (Fig. 1E) data was calculated based on trial-averaged evoked responses to 16 binocularly presented moving grating stimuli equally spaced between 0° and 360°. Orientation preference was computed by vector summation:

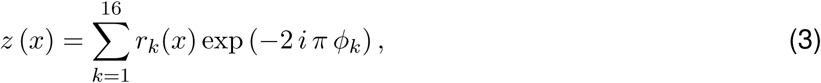

where *r*_*k*_(*x*) is the trial-averaged response to a moving grating with direction *φ*_*k*_ for pixel (widefield) or cell (two-photon) *x*. The preferred orientation was defined as *θ*(*x*) = 0.5 arg(*z*(*x*)) and orientation selectivity as abs 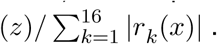

#### Intra-trial stability

To measure the reliability of the visual grating evoked responses within an individual trial (Fig. 1G, J), we compared the response pattern at two different time points *t*_1_ and *t*_2_ after stimulus onset (1 s vs. 3 s, respectively, for widefield and 0.5 s vs. 2.5 s for two photon imaging data). First, we computed the correlation

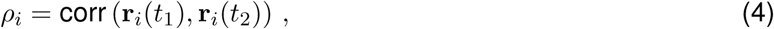

where **r**_*i*_(*t*_1_) and **r**_*i*_(*t*_2_) are the response patterns for the *i*-th trial at times *t*_1_ and *t*_2_, respectively. ’Intra-trial stability’ is defined by averaging this quantity over all trials from all grating directions (*N*_Evoked_ = *N*_stim._ × *N*_trial_ trials in total).

#### Dimensionality

We estimated the linear dimensionality *d*_eff_ of the subspace spanned by either evoked (Fig 2C, Fig 3E, Fig 4D, M) or spontaneous (Supplementary Fig. 3E) activity patterns by the participation ratio (*8*). To cross-validate our estimates of the spectrum of the covariance matrix, we split the response patterns from an imaging session into two equally large non-overlapping sets *A* and *B* (equal splits for each stimulus, individually), and used set *B* to estimate the principal component vectors and set *A* to estimate the variance for each of these components by applying Eq. (1) above. Our measure of ’dimensionality’ is then defined by

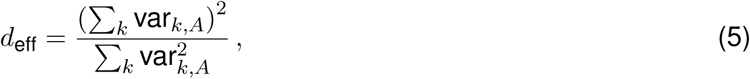

where var*_k,A_* is the cross-validated estimate of the k-th variance component. We repeated this calculation for *N* =100 different random splits into sets *A* and *B*. All values reported are averages over these *N* repetitions.

#### Alignment with spontaneous activity

To assess the overlap of evoked activity with the prevalent linear dimensions of spontaneous activity (on the same day; Figs 4F,G and 3G), we calculated the fraction of evoked pattern variance that is explained by the k-th principal component of the spontaneous pattern. To do so we used Equation (1), where *A* and *B* are the sets of evoked and spontaneous patterns, respectively.

The quantity ’alignment to spontaneous activity’ (Fig. 2H) was defined as the fraction of total sponta-neous pattern variance explained by an individual evoked pattern,

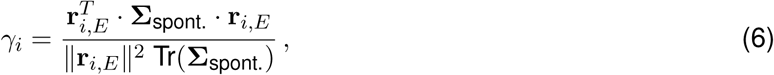

averaged over all *N*_evoked_ evoked patterns. Here, **r**_*i,E*_ is the *i*th evoked pattern taken 0.5 s after stimulus onset and **Σ**_spont_. the covariance matrix of the spontaneous patterns.

#### Tests of model predictions

Our model predicts that individual responses that are more aligned to spontaneous activity 0.5 s after stimulus onset also tend to show a higher value of intra-trial stability during the subsequent stimulation period (Fig. 4G). We tested whether the empirical data confirms this relationship by computing, for a given animal on a given day, the correlation

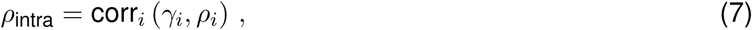

where *γ*_i_ is the alignment of an evoked pattern ri to spontaneous activity (Eq. 6), and *ρ*_i_ is the intra-trial stability of this evoked pattern (Eq. 4). As described above, the alignment to spontaneous activity was computed 0.5 s after stimulus onset, while the intra-trial stability was assessed by comparing the response between 1 s and 3 s after stimulus onset. The correlation *ρ*_intra_ is displayed in Fig. 4J.

Likewise, the model also predicts that individual stimuli, for which the average response is more aligned to spontaneous activity, also show higher values of trial-to-trial stability (SI Fig. 7A). To test this prediction we computed the correlation

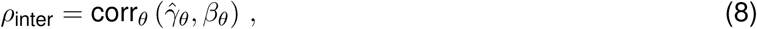

where 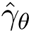 is the alignment of the trial averaged evoked pattern for stimulus *θ* to spontaneous activity (computed by replacing the response with its trial average in Eq. (6), and *β*_*j*_ is the trial-to-trial correlation for that stimulus (Eq. 2). The correlation *ρ*_inter_ is displayed in Fig. 4L.

Moreover, to test for a relationship between alignment to spontaneous activity and dimensionality, we assessed the degree to which these two quantities co-vary across the animals in a given age group (Fig. 4M, SI Fig. 7E). Here, we only used the first principal component **p**_1_ of the evoked pattern set to estimate alignment of evoked to spontaneous activity in order to avoid biases of dimensionality towards smaller numbers (as, for instance, the average alignment of individual patterns would potentially do). Concretely, we used Eq. (6), but with **r***_i,E_* being replaced by **p**_1_. We then assessed for each age group individually the correlation (across animals) between this alignment measure and the evoked dimensionality (Fig. 4M). Finally, by performing the analogous analyses for the trial-to-trial correlation (SI Fig. 7E, left) and intra-trial stability (SI Fig. 7E, middle) we also tested for the relationship between these quantities and the alignment to spontaneous activity on an animal by animal level.

### 2.4 Statistical analysis

We used re-sampling methods for hypothesis testing. Significance was assessed by comparing the average value of a given statistics between the real data and a suitable control distribution. The p-value was computed as the fraction of values in the control distribution that were more extreme than the difference observed in the real data.

#### Random control patterns

To test for the significance of a statistics characterizing the structure of a set of evoked or spontaneous patterns (for example, the trial-to-trial correlation in Fig. 1D), we computed the same statistics for *N* = 1000 control sets, each one of which comprises an equal number of surrogate patterns as the original set. The patterns in a control set exhibit the identical spatial correlation functions as the original patterns, but all statistical relationships beyond, i.e. higher order statistics and relationships among patterns, are eliminated. To generate one control set, we produced one surrogate pattern for each real pattern by computing its Fourier-transform, randomly shuffling the Fourier-phases (while preserving the amplitudes), back-transforming it into real space and re-applying the ROI mask of the real pattern. Repeating this procedure *N* times using different shuffles generated *N* control sets, from which we computed *N* control values of the statistics of interest, the distribution of which we compared with the real value.

#### Unpaired bootstrap test

To test whether a group average differs significantly between two groups (as for instance the difference between the response properties in normally reared vs. deprived animals in Fig. 3), we used an unpaired test based on a control distribution of group differences generated via bootstrap resampling. To generate control group differences, we pooled the data from the original two groups, drew two new groups of the same size from this pool by randomly drawing samples with replacement, and then computed the difference between the group averages. Repeating this procedure *N* = 1000 times resulted in a control distribution of group averages, which we compared with the real difference.

#### Paired permutation test

To test whether a group of animals showed a significant change across age (for instance, to establish whether the increase in trial-to-trial correlation in Fig. 1D is significant), we employed a paired permutation test to generate a control distribution. We randomly flipped the data points for a given animal between age groups and recomputed the difference between the group averages. Repeating this procedure *N* = 1000 times resulted in a control distribution, which we compared with the real difference.

Error bars were, if not mentioned otherwise, computed as the standard error of the mean (SEM) within a given age-group.

## 3 Network model

### 3.1 Motivation

We used a network model to study the feedforward-recurrent alignment hypothesis, which states that the experience-driven maturation of faithful visual representations after eye opening involves the proper alignment of visual input to the cortical recurrent network, such that this input gets sufficiently amplified through the network interactions. As a simple example for such amplification mechanism, consider an input pattern that primarily drives network units that also excite each other through their recurrent interactions. Such input is expected to elicit a response that is more robust against external and intrinsic sources of noise than an input pattern that drives a population with mixed (i.e. positive and negative) interactions. In the former case the input ’resonates’ with the network, while in the latter case the input gets dispersed through the network interactions.

According to this hypothesis, in the experienced cortex the inputs are well-aligned with the recurrent cortical network, whereas in the visually naive cortex, when patterned visual stimuli drive the visual system for the first time, the alignment of these novel inputs is only poorly developed. The transition from poorly aligned to well-aligned inputs is proposed to explain - via the change in network amplification that such improvement in alignment implies - the changes in the network properties we observed with the onset of visual experience, i.e. the development of reliable evoked responses that reside in a low-dimensional manifold, aligned with spontaneous activity. To explore this possibility, we studied a computational network model, which allowed us to cast this feedforward-recurrent alignment hypothesis into concrete mathematical terms, to test whether it can account for the changes in the network properties we observed during the maturation of visual representations and to derive further predictions that we tested using our empirical data.

### 3.2 Model setup

The model is intended to provide a concrete, yet as simple as possible formalization of this feedforward-recurrent alignment hypothesis in terms of a minimal linear network model of stimulus-evoked and sponta-neous neural population activity in layer 2/3 of the early visual cortex. We assumed that prior to eye opening a sufficiently strong recurrent network of excitatory and inhibitory neural populations is established in the visual cortex. This assumption is consistent with the pronounced, widespread, modular patterns of spontaneous activity seen in excitatory neural populations in the ferret visual cortex at this early stage in development, and the observation that such activity persists even after silencing the LGN (*2*). The assumption of strong inhibition, specifically, is consistent with the observation that these early spontaneous activity patterns in excitatory neural populations critically depend on network inhibition, which itself is organized into widespread modular patterns of activity that co-align with those in excitatory populations (*9*). Furthermore, we assumed that at eye opening, when structured visual stimuli drive the visual system for the first time, the relationship between the recurrent network previously established in layer 2/3 and this novel type of visually driven input is random. Moreover, visually driven input was assumed to be stochastic, reflecting various noise sources along the visual pathway including those within the cortex. Finally, we modeled spontaneous activity as the response of the recurrent network to an aligned, but relatively broad (i.e. high-dimensional) drive, assuming that spontaneous activity reflects inputs from a wide range of different sources.

We modeled the recurrent neural network using a standard firing rate equation

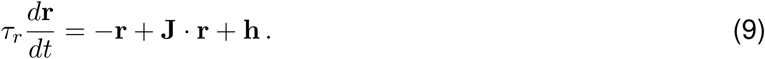

Here, ***r*** is the vector of rate units. Each unit *r*_*i*_, with *i* = 1, …, *n*, represents the pooled activity in a local group *i* of cortical neurons. The *n* × *n* matrix **J** describes the net interactions between these local pools of neurons, which can be positive or negative. To keep the network as generic and simple as possible, we assumed the network structure to be random and symmetric with interaction strengths drawn from a normal distribution *N* (*µ*_J_, *σ*_J_) with mean *µ*_J_ = 0 and standard deviation *σ*_J_ = *R*/2√*n*. Here, *R* describes the overall strength of recurrent interactions. In all figures we used *R* = 0.85, ensuring that the recurrent connections are strong enough to impact activity patterns. The size of the network was *n* = 200 (see Table 1 for a summary of parameters). The external (feedforward) input to the network is denoted by **h** = **h**_det_ +**h**_sto_ and consists of a deterministic part **h**_det_ and a stochastic part **h**_sto_, which are both assumed to be static, for simplicity, if not noted otherwise. Finally, we set the intrinsic timescale *τ*_*r*_ equal to 1 so all time is measured in units of this time constant.

### 3.3 Network amplification

Following standard procedures (e.g. (*10*)), the solutions to Eq. (9) are efficiently expressed in terms of the eigenvectors of the recurrent interaction matrix **J.** This matrix is real and symmetric and assumed to have full rank, hence all eigenvalues *λ*_*k*_ are real and the eigenvectors *e*_*k*_ form an orthonormal basis. In this basis, the dynamics decouples and the firing rate is given by

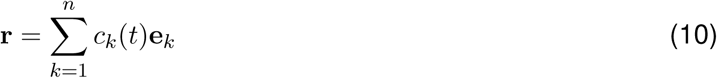

with coefficients *c*_*k*_ that depend on the overlap of the external input and the eigenvectors. For the connectivity defined above, the largest eigenvalue is approximately *λ*_max_ ≈ *R*. For an input constant in time, the steady-state solution of Equation (9) is given by

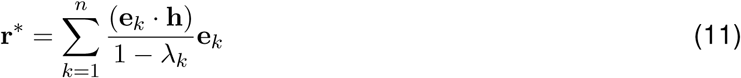

showing that the network amplifies an input stronger the more it overlaps with eigenvectors exhibiting a large eigenvalue. Consequently, the network responses tend to be biased towards the dimensions spanned by these leading eigenvectors, which can suppress unspecific noise components and result in activity distributions much lower-dimensional than the distribution of inputs, such as, for instance, in our model of spontaneous activity (Fig. 4E).

The alignment of a feedforward input **h** with the recurrent network **J** was defined as

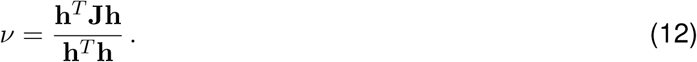

Rewriting the interaction matrix as

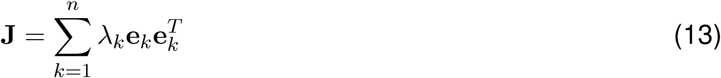

we observe that the maximally possible alignment is assumed when the input is proportional to the eigen-vector e_max_ with maximal eigenvalue *λ*_max_, in which case the alignment is *ν*_max_ = *λ*_max_ ≈ *R*. According to Eq. (11) such maximally aligned input also gets amplified the most by the recurrent network. A random input, instead, typically overlaps with a broad range of eigenvectors with mixed positive and negative eigenvalues and, therefore, its alignment will be close to zero and its net amplification much smaller. Note that such alignment-dependent network amplification is a generic property of recurrent networks and arises also in randomly structured networks, provided the recurrent interactions are sufficiently strong.

### 3.4 Population response properties

Our goal was to study whether the core network response properties considered in this study (i.e. inter- and intra-trial stability, dimensionality and alignment with spontaneous activity) improve substantially by aligning the feedforward input with the recurrent network structure (while keeping all other features of the input and the recurrent network unchanged). In the following we describe how we studied these response properties in the model.

#### Trial-to-trial correlation

To study the reliability of the network response to repeated presentations of the same stimulus in the model and to analyze how this reliability depends on feedforward-recurrent alignment (Fig. 4B), we assumed that the deterministic part of the stimulus-induced input to the network, det, is stimulus-dependent and fixed across different trials, while the stochastic part, **h**_sto_, varies from trial to trial and is stimulus-independent, for simplicity. The norm of the deterministic part of the input vector was set to 1, without loss of generality, see Table 2 for summarized parameters. Concretely, we considered inputs drawn from the multivariate normal distribution

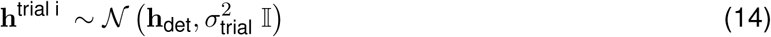

with stimulus-dependent mean **h**_det_, and diagonal covariance matrix with strength *σ*_trial_^2^ Given this input the distribution of evoked patterns is

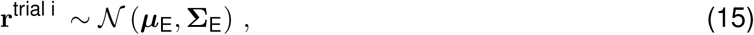

where (using Eq. 11) the mean response is given by

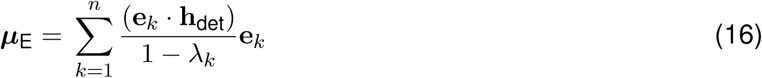

and the covariance structure by

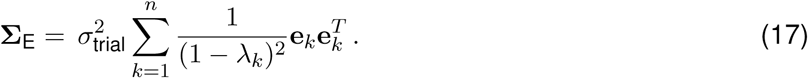

Analogously to the experimental data, we computed trial-to-trial correlation by drawing response vectors (N=100) from the pattern distribution (see Eq. 15) for a given input **h**_det_ and taking the mean over all possible pairwise correlations between these response vectors. Fig. 4B upper shows histograms over these pairwise correlations for a maximally aligned input, i.e. **h**_det_ equal to the eigenvector with largest eigenvalue *λ*_max_ (blue), and randomly aligned input of equal strength (green). To reveal the systematic dependency of trial-to-trial correlation on feedforward-recurrent alignment (Fig. 4B lower), we chose **h**_det_ proportional to the eigenvectors with ascending eigenvalues. Note that sampling the mean inputs from a broader distribution, such as the 10-dimensional Gaussian distribution defined in Dimensionality (see below) led to similar results.

#### Intra-trial stability

To study the stability of the network response to a sustained stimulus and how this stability depends on the alignment of the stimulus-induced input to the recurrent network (Fig. 4C), we assumed as input h a stationary Gaussian stochastic process that is uncorrelated in space and time with variance 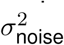, i.e. 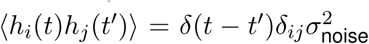 and time-independent mean **h**_det_ that depends on the stimulus. The stochastic differential equation for the external input is then d**h** = **h**_det_dt + *σ*_noise_ d**W**, where **W** is the Wiener process, and the stochastic differential equation for the network activity

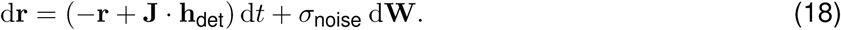

We numerically integrated the equation above for a duration of T = 120 with the steady-state solution hdet (see Eq. 11) as initial condition, using the Euler-Maruyama scheme with a time-step Δ*t* = 0.1 (see Table 3).

Analogously to the experimental data, we computed intra-trial stability as the correlation 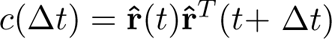 between z-scored response vectors 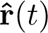 with time-lag Δ*t*. This correlation depends on the strength of the noise relative to the signal and the temporal correlations induced by the recurrent connections. As the temporal distance for assessing intra-trial stability in the experimental data is relatively large (2 s), we assumed that most temporal correlations induced by the recurrent network can be neglected and chose a sufficiently large time difference Δ*t* = 20. To obtain a more accurate estimate, we averaged *c*(Δ*t*) over all time points 0 ≤ *t* ≤ 100. Fig. 4C upper shows the normalized time-dependent response of the network unit with maximal time-averaged response, for a maximally aligned input (blue), and a randomly aligned input of equal strength (green). As above, to reveal the systematic dependency on feedforward-recurrent alignment, we chose hdet to match eigenvectors with ascending eigenvalues (Fig. 4C lower).

#### Dimensionality

To study in the model how the dimensionality of stimulus-evoked activity depends on feedforward-recurrent alignment (Fig. 4D), we applied time-independent inputs **h**^Dim^ to the network drawn from a low-dimensional multivariate Gaussian distribution **h**^Dim^ ∼ *N* (0, **Σ**^Dim^) with predefined dimensionality. As for the experimental data, dimensionality was defined by the participation ratio (*8*) *d*_eff_ = 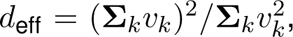, where *v*_*k*_ are the eigenvalues of the covariance matrix **Σ**^Dim^. To construct **Σ**^Dim^ we chose these eigenvalues proportional to an exponentially decaying function, i.e.

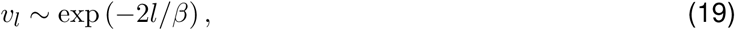

where *β* is a parameter. This models the decay of the variance explained by the leading principal components of evoked activity observed in our experimental data (see Fig. 2B) and yields a dimensionality *d*_eff_ ≈ *β*, which can be seen from the following argument: Inserting the expression (19) into the participa-tion ratio we obtain

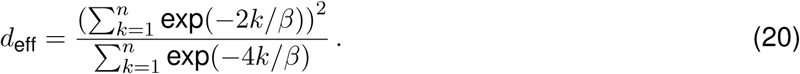

In the regime relevant to us 1 ≪ *β* ≪ n holds and thus we can approximate the sums by indefinite integrals, i.e.

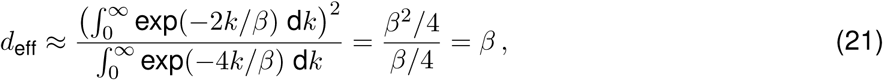

showing that in this regime the effective dimensionality is indeed approximated by *β*. In Fig. 4D, we used *β* = 10. We chose the following procedure to systematically vary the degree of alignment of this low-dimensional input to the recurrent network. We ordered the eigenvectors *e*_*k*_ of the interaction matrix **J** according to their eigenvalues *λ*_*k*_ in descending order, such that e_1_ is the eigenvector with the largest eigenvalue *λ*_max_ and e_*N*_ the eigenvector with the largest negative eigenvalue *λ*_min_. We then assigned variances with exponentially decaying magnitude (according to Eq. 19) to *M* = *κβ* < *n* of these vectors, starting with a leading eigenvector e_*L*_ and proceeding with the index in ascending order. The index *L* thus effectively determines alignment and the covariance matrix is then given by

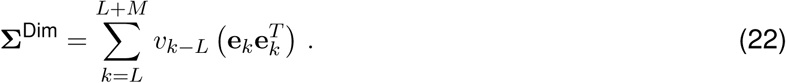

Truncating the sum at *κβ* results in an additional imprecision in the dimensionality, which can be estimated by performing the integrals in Eq. (21) only up to an upper bound *κβ*. This results in

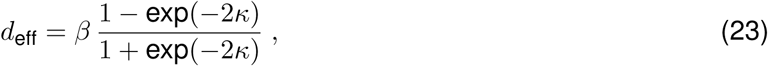

which is negligible for sufficiently large *κ*. We used *κ* = 5 in which case the truncation error is less than 0.01%.

To compute the activity spectrum for a maximally aligned input (Fig. 4D upper, blue line) we chose *L* = 1. For a random input (Fig. 4D, upper, green line) we used *n* orthonormal random vectors instead of the eigenvectors. To systematically study how dimensionality depends on feedforward-recurrent alignment (Fig. 4D lower), we varied L between 1 and *n*/2.

Given such low-dimensional Gaussian input distribution, the evoked activity patterns are then samples from a Gaussian distribution with covariance structure

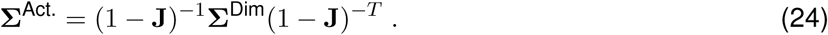

Because the input covariance **Σ**^Dim^ and the interaction matrix J share an eigenbasis per construction, the eigenvalues of **Σ**^Act^. can be expressed as

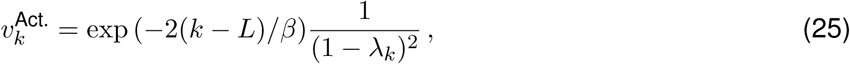

for *k* = *L*, …, *L* + *M* . The dimensionality of the evoked activity space can then be expressed as

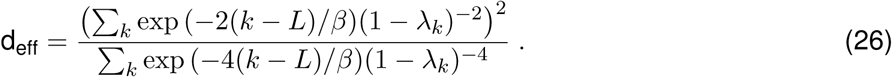

#### Alignment with spontaneous activity

We assumed spontaneous activity reflects inputs from a wide range of different sources, and so we modeled these inputs as samples drawn from a broad Gaussian distribution (Fig. 4E). Moreover, given that spontaneous activity is present more than a week prior to eye opening, we assumed this distribution is already aligned to the recurrent network at eye opening and remains aligned, subsequently. We modeled this distribution as

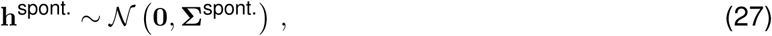

where **Σ**^spont^ is defined as **Σ**^Dim^ (Eq. 22), but using *L* = 1 and *β*_spont_. = 20. Spontaneous activity patterns are modeled as steady-state responses to these inputs and, following Eq. (24), are given by samples from the distribution **r**_*i,S*_ ∼ *N* (0, **Σ**_s.act._), where the covariance structure is given by

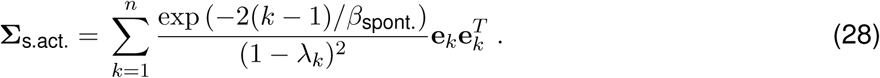

Analogous to the empirical data, we computed the alignment of evoked patterns with spontaneous activity using Eq. (6).

### 3.5 Network parameters

See Tables 1, 2 and 3 for a summary of parameters.

**Table 1:** Network parameters

| Parameter | Symbol | Value |
| --- | --- | --- |
| Number of network neurons | $n$ | 200 |
| Mean of weight distribution | $\mu_J$ | 0 |
| Magnitude of maximal eigenvalue | $R$ | 0.85 |

**Table 2:** Simulation parameters

|  |  |  |
| --- | --- | --- |
| Number of trials | $N_{\text{trial}}$ | 100 |
| Noise strength (trials) | $\sigma_{\text{trial}}$ | 0.02 |
| Noise strength (time) | $\sigma_{\text{time}}$ | 0.3 |
| Evoked dimensionality | $\beta_{\text{evoked}}$ | 10 |
| Spont. dimensionality | $\beta_{\text{spont.}}$ | 20 |

**Table 3:**
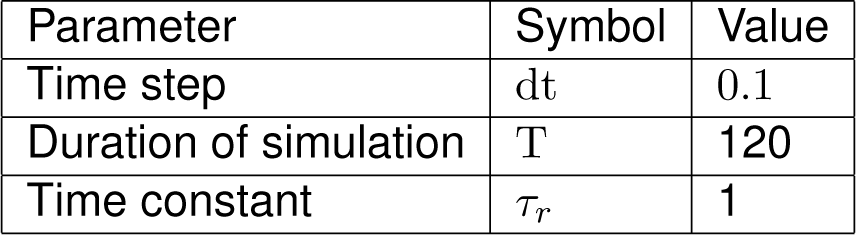
Numerical parameters

All networks were simulated using custom-written code in python.

## Supplementary Information

**SI Figure 1:**
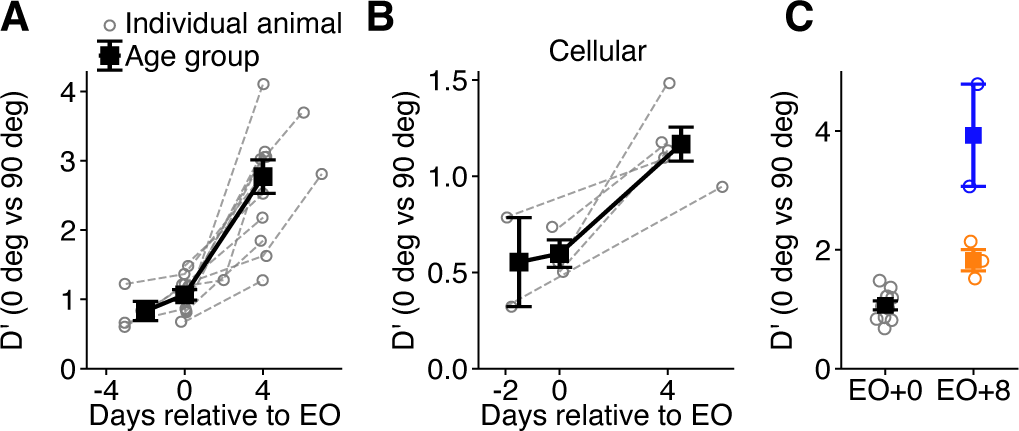
The strong but highly variable network responses in visually naïve and deprived animals are accompanied by relatively weak orientation discriminability. (Related to Figs. 1 and 3) (A) Discriminability index D’ (Cohen’s d) for distinguishing responses for a 0 and 90 deg grating stimulus. Data obtained from the same animals as in Fig. 1C. Error bars show Mean ± SEM for age groups (EO-3 to −2, EO+0, EO+4 to +7, N= 4,11,11 animals, respectively. (B) Same as (A), but for cellular responses. Same animals as in Fig. 1I. (C) Same as (A) but for deprived (N=3, orange) and control (N=2, blue) animals, in comparison to visually naïve animals at EO (black, reproduced from (A)). Circles: single animals; error bars: Mean ± SEM. Same animals as in Fig. 3C.

**SI Figure 2:**
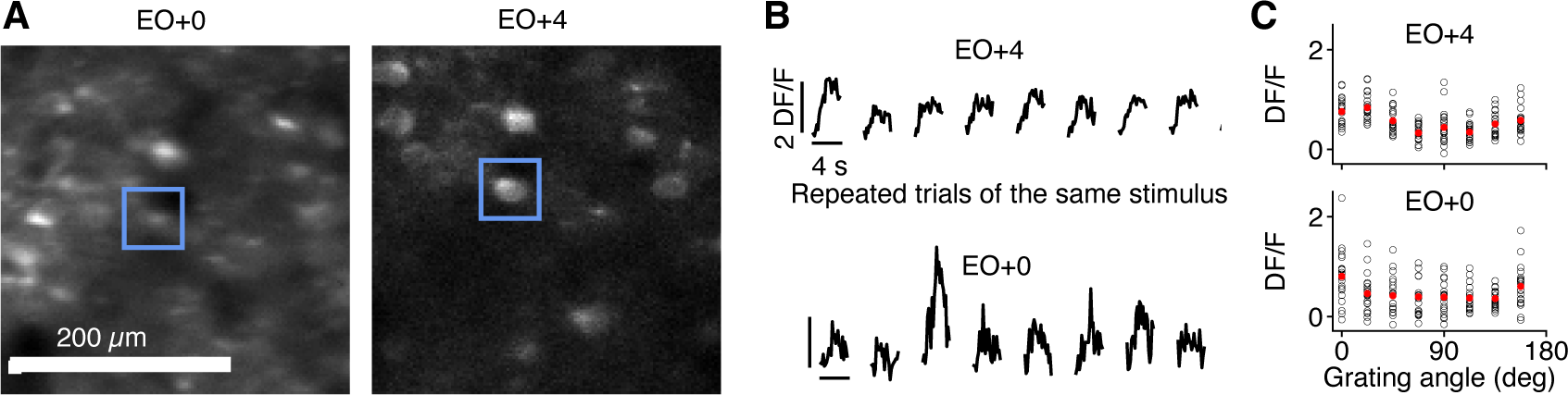
Chronic two photon calcium imaging of visual grating evoked activity in the early developing visual cortex reveals a high degree of response variability in visually naive animals in populations of individual cells. (Related to Fig. 1H-J) (A) Representative example of chronic two photon calcium recordings in the early developing ferret visual cortex (used in Fig. 1H-J). Approximately the same cortical region was imaged in a visually naive cortex (left) and after four days with visual experience (right). For illustration purposes, only about one quarter of the total imaging field of view is depicted. A cell tracked over the two time points is marked by the blue box. (B) Visual grating evoked calcium traces (same stimulus, six trials) for a representative cell (marked in (A) by the blue box) in the visually naïve (bottom) and experienced (top) cortex. (C) Single trial responses (averaged over the entire stimulus period of 4 s) of the same cell as in (B) as a function of grating orientation. Open black circles: individual trials; solid red circles: trial averages.

**SI Figure 3:**
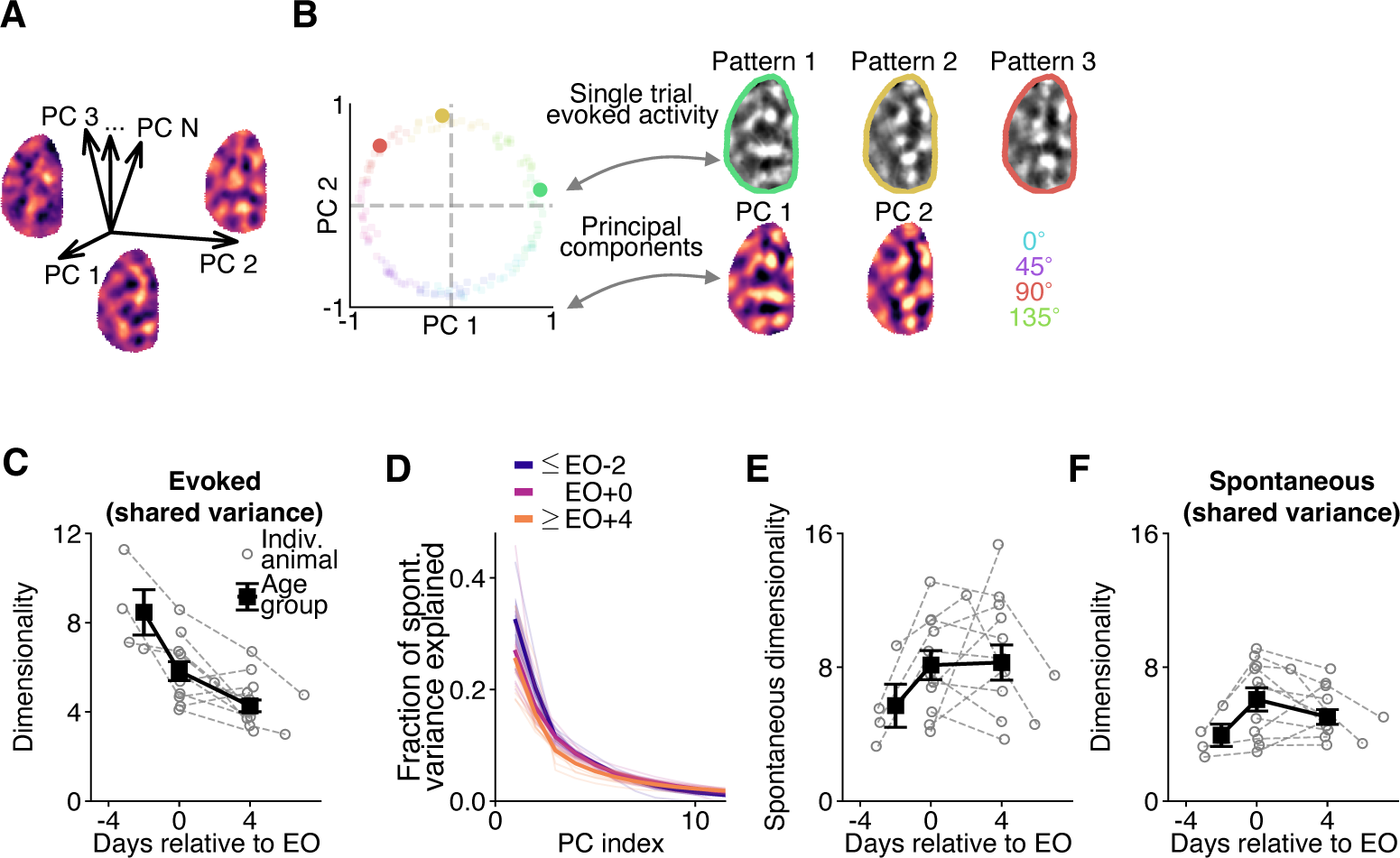
Additional analyses regarding the dimensionality of visual grating evoked and spontaneous activity. (Related to Fig. 2B-E) (A) Activity patterns can be expressed as a weighted superposition of principal components (PC), which themselves can be represented as patterns covering the same cortical ROI as the activity patterns. Three example PC are illustrated in red. (B) Large bold circles: three examples of evoked patterns projected into the space spanned by the first two PC of evoked activity. These patterns and the first two PCs are illustrated (in grayscale and red, respectively). Small faint circles: Projections of all evoked patterns (reproduced from Figure 2B). (C) Dimensionality estimates based on shared variances across two non-overlapping subregions in the ROI (following the method in Stringer et al., 2019) yield an age trend similar to our estimates in Fig. 2C. To define the two subregions, we divided the ROI into N nonoverlapping stripes of width 392 μm and assigned stripes with an odd index to subregion 1 and those with an even index to subregion 2. Pre-processing of calcium activity patterns was applied separately in these two subregions. (D) Fraction of variance of spontaneous activity explained by its leading PC for different age groups (EO-3 to −2, EO+0, EO+4, EO+6: N= 4,11,11,2 animals, respectively; thick lines, average over animals; thin lines, individual animals; see Methods). (E) Linear dimensionality of spontaneous activity across age (participation ratio, cross-validated, see Methods). (F) As (B), but for spontaneous activity. Note that the number of patterns is matched across age, but not between evoked and spontaneous activity, hindering a direct comparison of their dimensionality.

**SI Figure 4:**
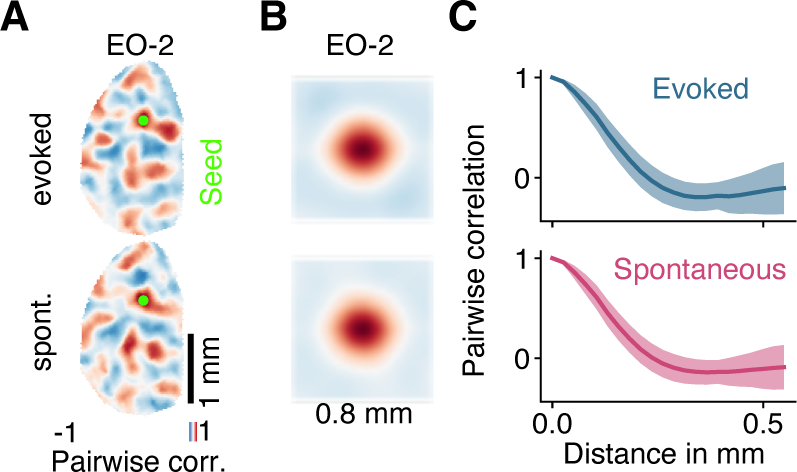
At a local, modular scale, the initial grating evoked patterns and the patterns of the endogenous network exhibit a similar correlation function. (Related to Fig. 2) (A) Example correlation patterns (Smith et al., 2018) for total signal correlation (top row) and spontaneous correlation (bottom row) for the same seed point in a visually naïve ferret. Scalebar: 1 mm. (B) 2D correlation function of evoked (top) and spontaneous activity (bottom) at EO-2. Correlation patterns from (A, left) were averaged over all possible seed-points, displayed for a diameter of 0.8 mm. (C) Radial profiles of correlation functions from (B) with SD over seed-points. Note the similar location and depth of the trough, indicating a similar spatial scale and degree of order in the local modular structure of the spontaneous and initial evoked activity.

**SI Figure 5:**
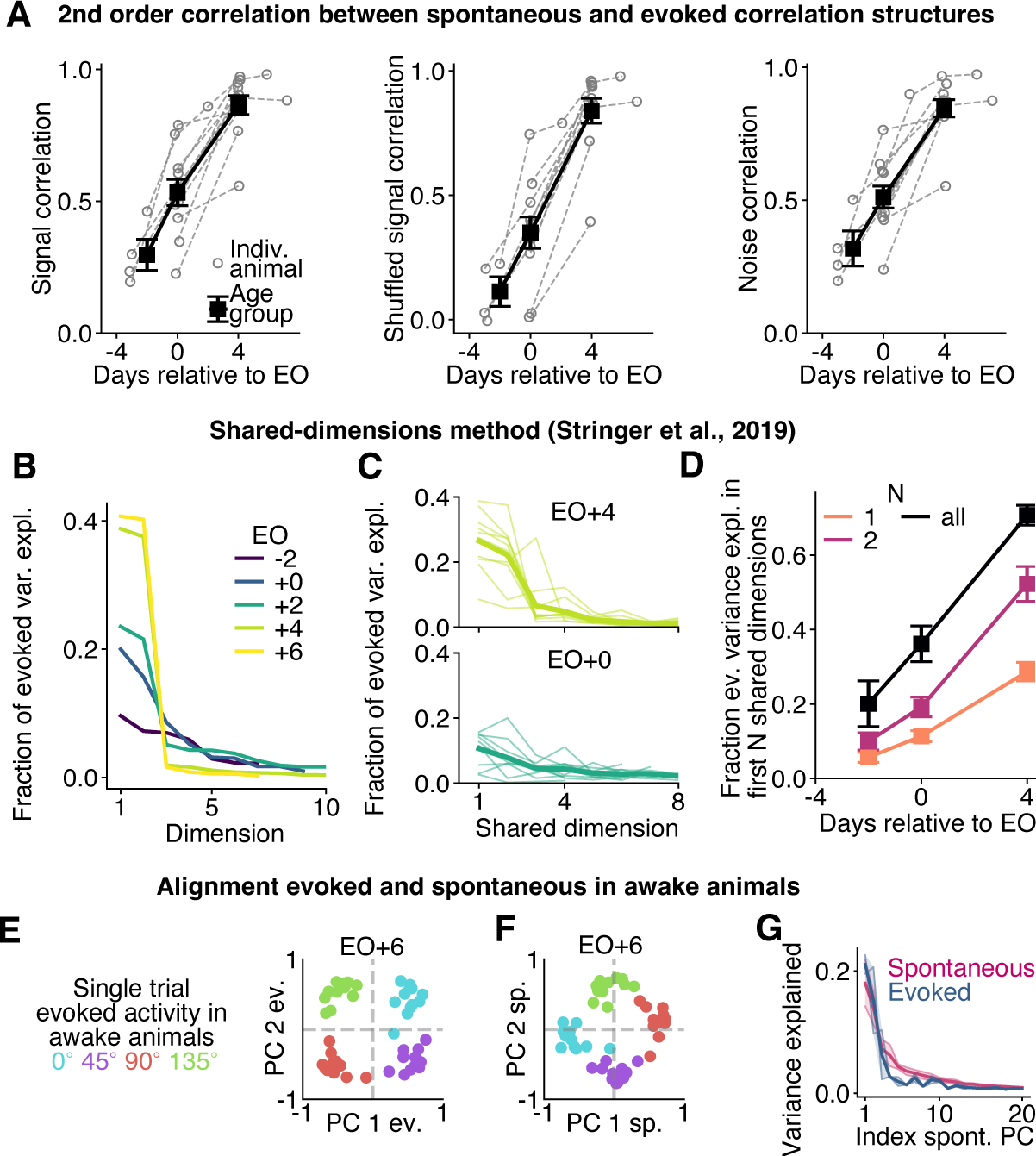
Additional analyses of the alignment of spontaneous and visual grating evoked activity. (Related to Fig. 2F-H) (A) As a first alternative to the analysis in Fig. 2F-H we computed the 2^nd^ order correlation between the spontaneous correlation and several evoked correlation structures, namely the total signal correlation (left), the trial-split signal correlation (middle) and the noise correlation (right). Animals were longitudinally recorded. Same animals and age bins as in Fig. 1 D, G: (EO-3 to −2, EO+0, EO+4 to +7, N= 4,11,11 animals, respectively). Correlations calculated for the subset of the 20% most selective locations at EO+4 excluding pairs with a distance smaller than 465 microns. (B-D): As a second alternative to the analysis in Fig. 2F-H, we analyzed the shared dimensions between spontaneous and evoked activity (Stringer et al, 2019). (B) Fraction of variance explained by shared dimensions between spontaneous and evoked activity across age for the animal from Fig. 2D-G. Shared dimensions were calculated as linear combinations of the leading principal components of spontaneous activity (of the same age) accounting for 75% of its own variance. (C) Fraction of evoked variance explained by shared dimensions four days after EO (top) and at EO (bottom). Thin lines: individual animals; Broad lines: average over animals. N=11 animals. (D) Total fraction of evoked variance explained by N=1, 2, all (orange, magenta, black) shared dimensions, averaged over animals (N=11). Error bars: Group average and SEM. (E-G) Alignment in awake animals. (E) Projections of evoked patterns into the space spanned by the first two evoked PC, as in Fig. 2A top, but for an awake animal. (F) Projections of evoked patterns into the space spanned by the first two PC of spontaneous activity, as in Fig. 2G top, but for an awake animal. (G) Alignment of evoked to spontaneous activity in the awake cortex (N=2 animals; compare Fig. 2H). Data in (E-G) from Smith et al., 2018.

**SI Figure 6:**
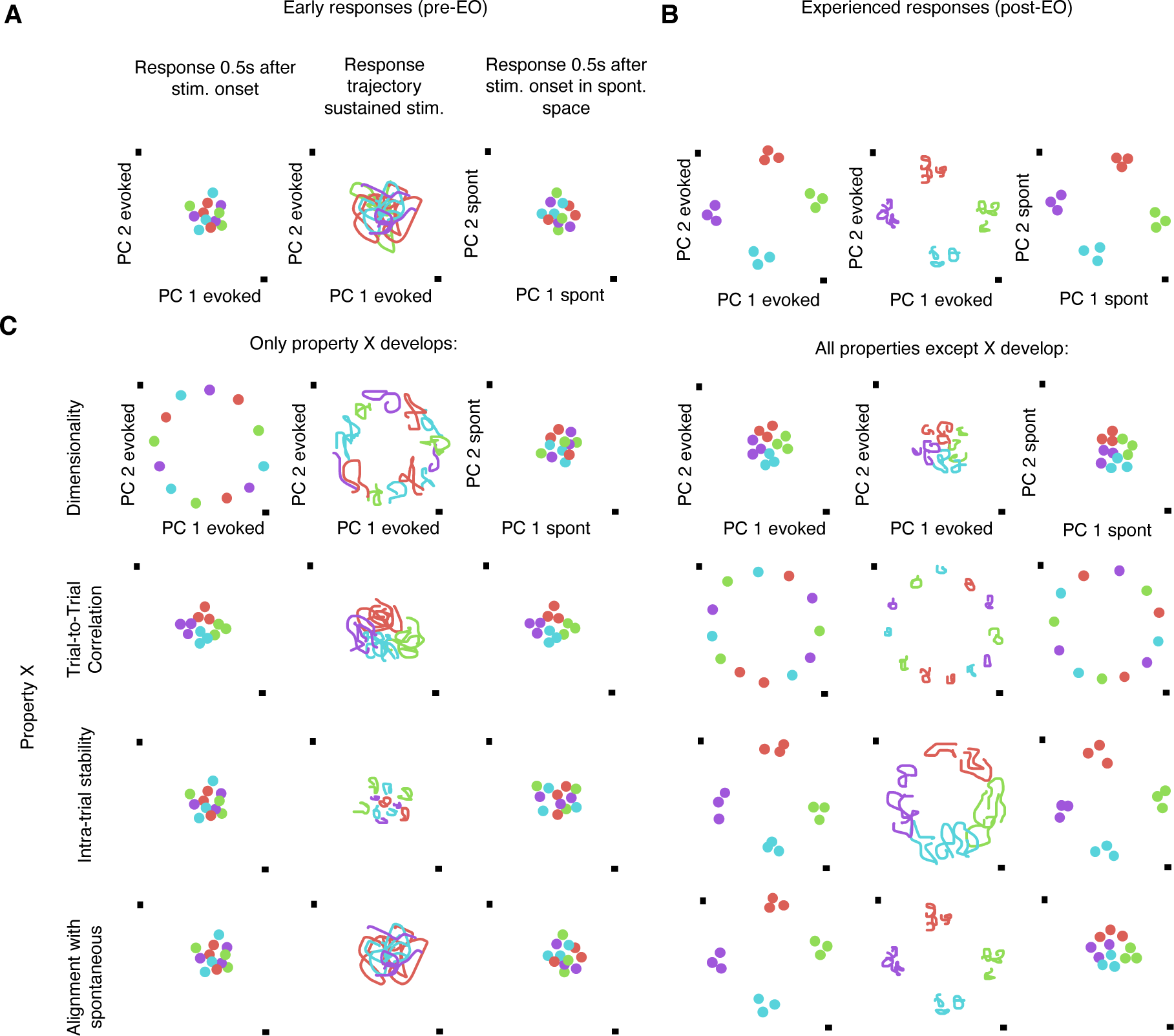
Network response properties can, in principle, change largely independently from one another. (Related to Fig. 4) Schematics of evoked activity projected onto the principal components (PC) of either evoked (compare Fig. 2A) or spontaneous activity (compare Fig. 2F, G) showcasing the four network properties in focus: trial-to-trial correlation (larger the smaller the scatter of responses with the same preferred orientation (marked by color)), dimensionality (smaller the larger the radius of projections onto evoked PCs), Intra-trial stability (larger the smaller the extend of trajectories of temporally evolving activity) and alignment with spontaneous activity (larger the larger the radius of projections onto spontaneous PC). (A) Evoked responses in visually naïve animals are relatively high-dimensional and show poor trial-to-trial correlation, intra-trial stability and alignment with spontaneous activity. (B) Evoked responses in experienced animals are low-dimensional and show high trial-to-trial correlation, intra-trial stability and alignment with spontaneous activity. (C) These four properties can develop largely independently from one another. The sketches show the schematics of evoked activity if only one of the four properties develops (left) and if all but one develops (right).

**SI Figure 7:**
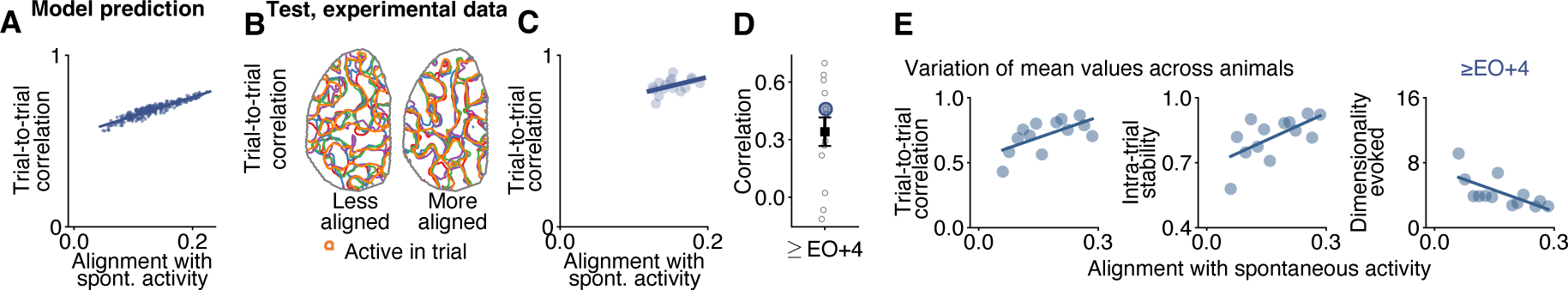
The model predicts a correlation between the alignment of the trial-averaged evoked pattern to spontaneous activity and trial-to-trial correlation, consistent with the empirical data. (Related to Fig. 4G-J) (A) Model prediction: single-trial responses to a stimulus for which the trial-averaged pattern is more aligned to spontaneous activity also show higher trial-to-trial correlation. N=300 samples drawn for the input distribution used in Fig. 4D with optimal alignment and added noise as in Fig. 4B. (B-D): Test in experiment. (B) Reproduced from Fig. 4K. Contours show outlines of grating evoked patterns (taken 0.5 s after stimulus onset) from different trials for a stimulus for which the average pattern is less (left) or more (right) aligned to spontaneous activity. Trial-to-trial correlation: 0.75 (left), 0.84 (right); alignment to spontaneous activity: 0.13 (left), 0.18 (right). (C) As (A) but for experimental data using all stimuli from the animal shown in (B) at EO+4. (D) Reproduced from Fig. 4L. Correlation for the data shown in (C) (large blue circle) and for all other animals in the oldest age group (grey circles; 13 experiments from N=11 animals at EO+4 to +7). Error bars show Mean ± SEM. respectively). (E) Trial-to-trial correlation, intra-trial stability and evoked dimensionality co-vary with the alignment of the first evoked PC to spontaneous activity (see Methods). 13 experiments from N=11 animals at EO+4 to +7. Parts of data reproduced from Figure 4M.

